# MACF1 Facilitates SMAD7 Nuclear Translocation to Drive Bone Formation in Mice

**DOI:** 10.1101/743930

**Authors:** Fan Zhao, Xiaoli Ma, Wuxia Qiu, Pai Wang, Ru Zhang, Zhihao Chen, Peihong Su, Yan Zhang, Dijie Li, Jianhua Ma, Chaofei Yang, Lei Chen, Chong Yin, Ye Tian, Lifang Hu, Yu Li, Ge Zhang, Xiaoyang Wu, Airong Qian

## Abstract

MACF1 is a large crosslinker that contributes to cytoskeleton integrity and cell differentiation. Loss of MACF1 impairs multiple cellular functions in neuron development and epidermal migration, and is the molecular basis for many diseases such as heart failure and Parkinson’s disease. MACF1 is highly abundant in bones, however, its involvements in osteogenic differentiation and bone formation are still unknown. In this study, by conditional gene targeting to delete the *Macf1* gene specifically in MSCs, we observed ossification retardation and bone loss in MACF1 deficient mice in different developmental stages, which we traced to disorganized cytoskeleton and decreased osteogenic differentiation capability in MSCs. Further, we show that MACF1 interacts and facilitates SMAD7 nuclear translocation to initiate downstream transcription. These findings are hopefully to expand the biological scope of MACF1 in bones, and provide experimental basis for targeting MACF1 in degenerative bone diseases such as osteoporosis.

## INTRODUCTION

### The microtubule-actin cross-linking factor 1 (MACF1) is an important mammalian spectraplakin protein that interacts with the cytoskeleton (Ka et al., 2014; Leung et al., 1999; Suozzi et al., 2012)

As one of the largest mammalian protein (∼ 620 KD) that coordinates the cytoskeleton, MACF1 has the ability to bind to actin filaments with its N-terminal calponin homology domain, and interact with microtubules with the C-terminal GAR/GSRA domain (Lane et al., 2017; Leung et al., 1999) (Figure EV1A). MACF1 also has cation binding sites in the C-terminus (Figure EV1A), and homologous sequence similar to the Smc family of ATPases (Lane et al., 2017; Wu et al., 2008), suggesting considerable potential kinase activity. In addition, MACF1’s cytoskeletal crosslinking capability together with its ATPase activity enable growth of the MTs toward focal adhesions (FAs), which facilitates cell migration and polarization (Wu et al., 2008). MACF1 mainly localizes in the cytoplasm, and is reported to participates in MT-mediated cargo transport (Chen et al., 2006; Kakinuma et al., 2004; Wu et al., 2008). MACF1 is also found in the nucleus, a few MACF1 goes to the nucleus from the cytoplasm.

MACF1 is expressed ubiquitously as early as at E7.5 during embryonic period (Chen et al., 2006), and plays essential roles in regulating various development processes in mammals (Antonellis et al., 2014; Ka and Kim, 2016; May-Simera et al., 2016). With especially high abundance in the heart, lung, brain and bones, its unique structure enables MACF1 a variety of functions in neuron development, epidermal migration and more (Duhamel et al., 2018; Escobar-Aguirre et al., 2017; Yue et al., 2016), analysis of these evidences show that MACF1 is crucial for normal cells to proliferate, polarize and differentiate. Reduction or depletion of MACF1 disorganizes the cytoskeleton and impairs cellular functions (Escobar-Aguirre et al., 2017; Ka et al., 2016; Ma et al., 2017; Wang et al., 2017), and further confers risk for degenerative diseases such as inflammatory colitis and Parkinson’s disease (Ka and Kim, 2016; Kodama et al., 2003; Ma et al., 2017; Wang et al., 2017).

### However, the function of MACF1 is less understood in bones

The bones are rigid structural support and endocrine organ in mammals. During early embryonic development, osteoprogenitors aggregate and differentiate into osteoblasts to synthesize new bone matrix and form flat bones such as the skull, while replacement of the cartilage tissue by ossified bone forms long bones such as the femur. In adults, bone metabolism is dynamicly regulated by osteoblasts-mediated bone formation and osteoclasts-mediated bone resorption (Khurana, 2009). MACF1 is also highly abundant in the bone tissue, but its functions in the regulation of bone development and formation remains unknown. Considering the vital roles MACF1 play in other tissues, the lack of information on MACF1’s function in bone tissue needs to be addressed, we wonder how MACF1 regulates bone development and formation. In addition, as a key cytoskeletal regulator, we wonder in particular how MACF1 affects the cytoskeleton in bone-forming cells and controls osteogenic differentiation. Our earlier studies show that knockdown of MACF1 in preosteoblasts inhibited osteoblast differentiation (Hu et al., 2017, 2015), and femoral *Macf1* level is significantly lower in osteoporotic mice (Yin et al., 2018) and is also negatively correlated to the age of mice (Hu et al., 2017), suggesting a close relationship between MACF1 and osteoporosis. Therefore, we hypothesize that MACF1 is essential for maintaining the cytoskeleton in mesenchymal stem cells (MSCs), and positively regulates osteogenic differentiation and bone formation.

### In this study, we unveiled essential roles of MACF1 in osteogenic differentiation and bone formation *in vivo*

By genetical manipulation to delete MACF1 in the mesenchyme, we identified essential roles of MACF1 in regulating osteogenic differentiation and bone formation in mice. We show that MACF1 is essential for early stage bone development, loss of MACF1 in MSCs retards bone development and further impairs bone properties and bone strength. *In vitro* experiments demostrate that MACF1 is essential for maintaining the cytoskeleton in differentiating MSCs, loss of MACF1 disorganizes the cytoskeleton and inhibit osteogenic differentiation. In addition, we show that MACF1 directly interacts with SMAD7 and facilitates SMAD7 nuclear translocation to initiate downstream osteogenic pathways. By coupling the results obtained, we have identified key roles of MACF1 in maintaining osteogenic differentiation and bone formation, and advanced the potential for exploiting MACF1 as a new therapeutic target for degenerative bone diseases such as osteoporosis.

## Results

### Mesenchymal deletion of MACF1

MACF1 is a large structural protein (Bernier et al., 1996; Leung et al., 1999). Previous studies show that conventional inactivation of the *Macf1* gene leads to teratogenicity and lethality during development (Chen et al., 2006), suggesting MACF1 is crucial for embryo development in mice. In order to study the *in vivo* involvement of MACF1 in bone development and formation, we utilized the Cre/Loxp technology and generated a genetically modified mouse model in which the *Macf1* gene was specifically deleted in mesenchymal stem cells (Flox, Macf1^f/f^; cKO, Macf1^f/f^;Prx1^Cre/+^) (Figure EV1A and S1B). Similar to those conditional MACF1 deficient mice generated using other Cre lines, mesenchymal deletion of MACF1 did not significantly affect body length or body weight in neonates (Figure 1A, Figure EV1C) (Goryunov et al., 2010), and offsprings were obtained in an expected sex ratio and Mendelian frequency (Figure EV1D), By adulthood, body size and organ coefficient remained unchanged as well (Figure 1B, S1E and S1F), suggesting that mesenchymal deletion of MACF1 did not apparently change fundamental physiological indications in mice.

**Fig. 1.**
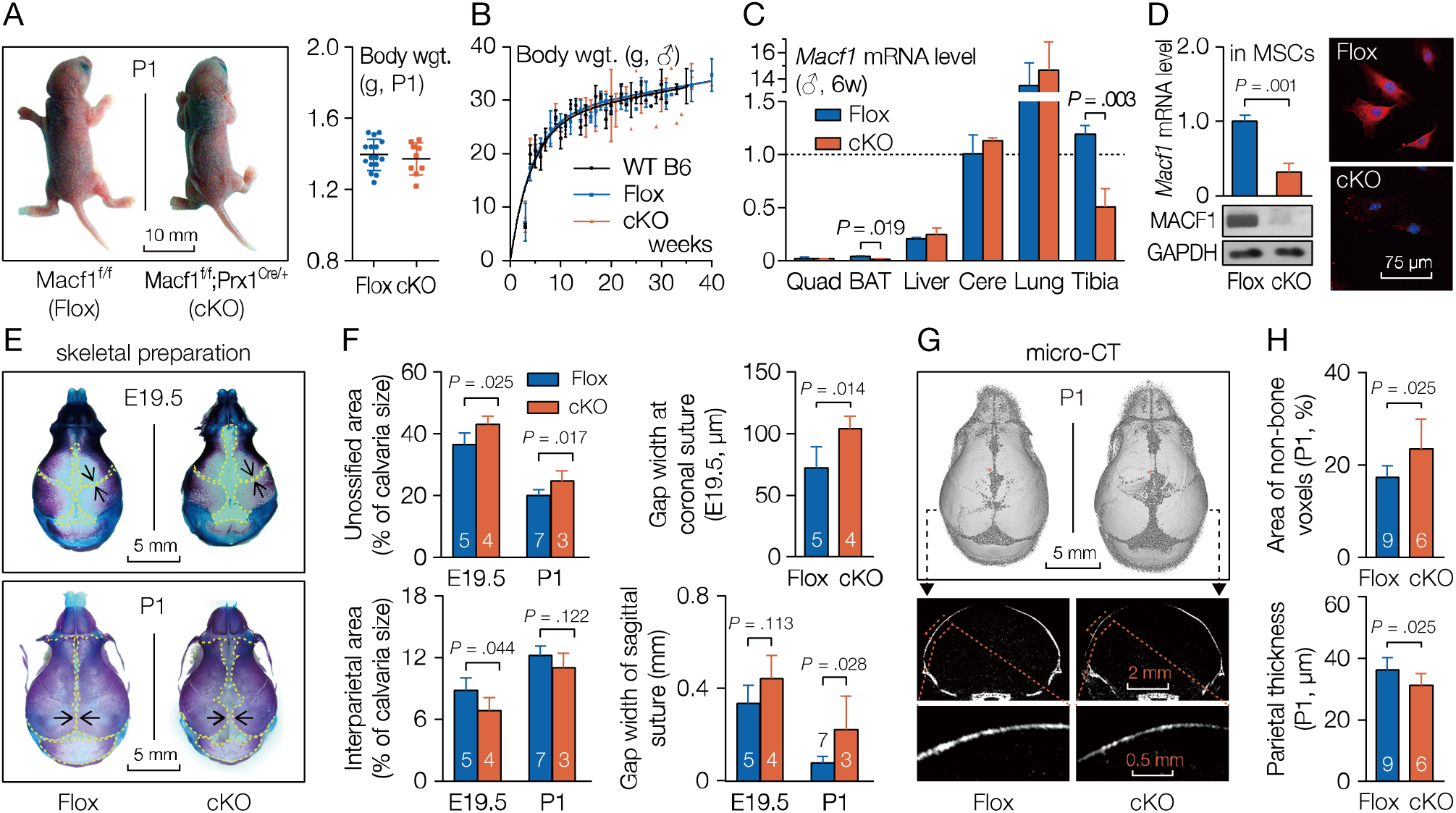
Mesenchymal deletion of MACF1 retards early stage bone development in mice. (A) Representative images and body weight of neonatal MACF1 cKO mice (P1, postnatal day 1). (B) Growth curve of male MACF1 cKO mice. Least-squares fitting Gompertz function was used. (C) Real-time quantitative PCR analysis of *Macf1* mRNA levels in organs or tissues of MACF1 cKO mice (6-week old, male). Quad, quadriceps. BAT, brown adipose tissue. Cere, cerebrum. (D) Expression and distribution of MACF1 in MACF1 deficient MSCs. *Gapdh*/GAPDH were used as internal reference. (E) Representative images of cleared skeletal preparation showing early-stage development of the skull (E19.5, embryonic day 19.5). Red shows ossified tissue, while blue indicates cartilage (dashed lines). (F) Quantification of panel E. Gap width indicates the nearest distance between adjacent bones. (G) Representative 3D reconstruction images showing skull development at P1. Cross-sections of the parietal bone are also shown. (H) Quantification of panel G; Non-bone voxels indicate unossified part. Data are represented as mean ± s.d. Significances were determined using student’s *t*-test.

We then detected MACF1 expression levels to confirm successful deletion of MACF1. Real-time quantitative PCR (qPCR) assays for detection of tissue *Macf1* levels showed that in the conditional knockout (cKO) group, the *Macf1* gene level was significantly lower in bone tissue, while in other tissues like muscle, cerebrum and liver, *Macf1* mRNA levels remained unchanged compared with control mice (Figure 1D). It may be worth noting that mRNA level of the *Macf1* gene was also lower in the brown adipose tissue (Figure 1D). Then immunoblotting and immunofluorescence experiments further demonstrated that MACF1 protein was barely detectable in MACF1 cKO MSCs (Figure 1E). The results indicate that conditional targeting of MACF1 was successful, and that mesenchymal deletion of MACF1 does not change fundamental physiological indications in mice.

### Mesenchymal deletion of MACF1 impairs early-stage bone development in mice

Previous studies report that Prx1-driven (paired related homeobox 1) Cre recombinase is expressed in a subset of craniofacial mesenchyme and in early limb bud mesenchyme (Elefteriou and Yang, 2011; Logan et al., 2002). To study the impact of MACF1 on bone development in mice, we first examined skull development and morphology at late embryonic stage as well as in neonatal mice. Cleared skeletal preparation of the skull showed that the cKO neonates displayed obvious retardation in cranial bone development at E19.5 and P1, manifested as delays in suture fusion and calvarial ossification (Figure 1E, 1F and S2A). MicroCT experiment again substantiated the retardation in suture closure, and found that parietal bone was thinner in cKO mice at P1 (Figure 2C and 2D). In addition, formalin-fixed paraffin embedded (FFPE) skull sections were also prepared. HE staining confirmed attenuation in calvaria thickness, while alkaline phosphatase (ALP) staining showed that cKO mice had less ALP expression in parietal bone (Figure 2E), but Alcian blue staining didn’t show significant difference between Flox and cKO mice (Figure EV2E), suggesting decreased osteoblastic activity in MACF1 deficient mice during cranial bone development.

**Fig. 2.**
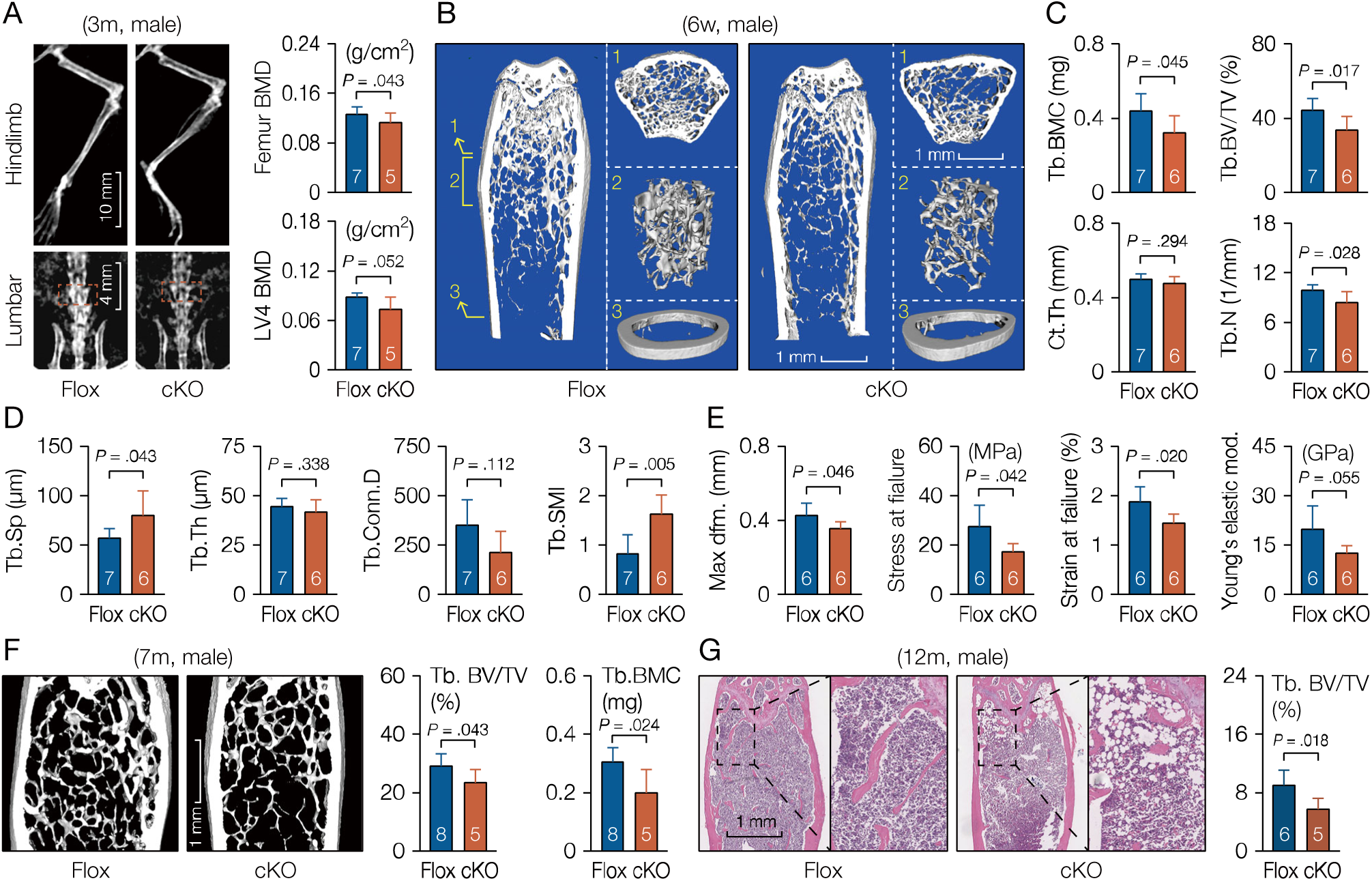
Mesenchymal deletion of MACF1 weakens bone properties in adult mice. (A) X-ray images of MACF1 cKO mice (male, 3-month old). BMD of femur and third lumbar vertebra were quantified. (B) Representative 3D reconstruction images showing microarchitecture of distal femur (male, 6-week old). Positions of reconstructed region are indicated by Arabic numerals. (C and D) Stereological parameters for trabecular and cortical bone in distal femur. BMC, bone mineral content; BV/TV, bone volume fraction; Ct.Th, cortical thickness; Ct.Ar, cortical area. Tb.N, trabecular number; Tb.Sp, trabecular spacing; Tb.Th, trabecular thickness; SMI, structural model index. Conn.D, connectivity density. (E) Femoral mechanical property (male, 3-month old). Max dfm., maximum deformation; Young’s mod., Young’s elastic modulus. (F) MicroCT analysis of distal femur (male, 7-month old). (G) Representative images of H&E staining showing trabecular in distal femur (12-month old, male, coronal sections). Bone volume fraction was quantified. Data are represented as mean ± s.d. Statistical significance were determined using student’s *t*-test.

Unlike flat bones, bones of the trunk and extremities are formed with an initial hyaline cartilage. To further investigate the effect of MACF1, long bones were also scrutinized. Cleared bone preparation for long bones revealed that mineralized bone formation in tibia was also slower due to MACF1 deletion (Figure EV2C and S2D), while other long bones such as ribs and forelimbs were less affected (Figure EV2C and S2D), indicating that, to some extent, development and formation of the lone bones in neonatal mice were also affected due to MACF1 deletion. The results suggest that MACF1 is crucial for early-stage development and formation of the skeleton in mice.

### Loss of MACF1 weakens bone properties in adult mice

Bone forming cells derived from mesenchymal stem cells are responsible for bone matrix synthesis and mineralization (Hilton, 2014), and finally form the mineral structure that supports bone functions. To investigate the involvement of MACF1 on bone phenotype during adulthood, bone properties in MACF1 deficient mice were tested. Firstly, 3-month old mice were subjected to dual energy X-ray absorptiometry (DEXA) scanning, the results showed that mineral density was lower in both femur and third lumbar vertebra (Figure 2A), bone mineral content and bone volume were also lower (Figure EV3A).

Next, distal femur from 6-week old male mice were scanned and reconstructed by microCT. The results showed that in cKO mice, bone mineral content and mineral density were lower (Figure 2B and 2C), manifest as decreased trabecular number, thickness and increased trabecular spacing (Figure 3D), indicating that MACF1 deletion decreased bone mass, and degenerated bone microarchitecture. In addition, to known mechanical properties of the bone, freshly-isolated femora were subjected to 3-point bending test. We found in cKO group not only reduced maximum compression stress and compression strain, but also reduced Young’s elastic modulus (Figure 3E), suggesting that toughness and strength of the bone were weakened in cKO mice.

**Fig. 3.**
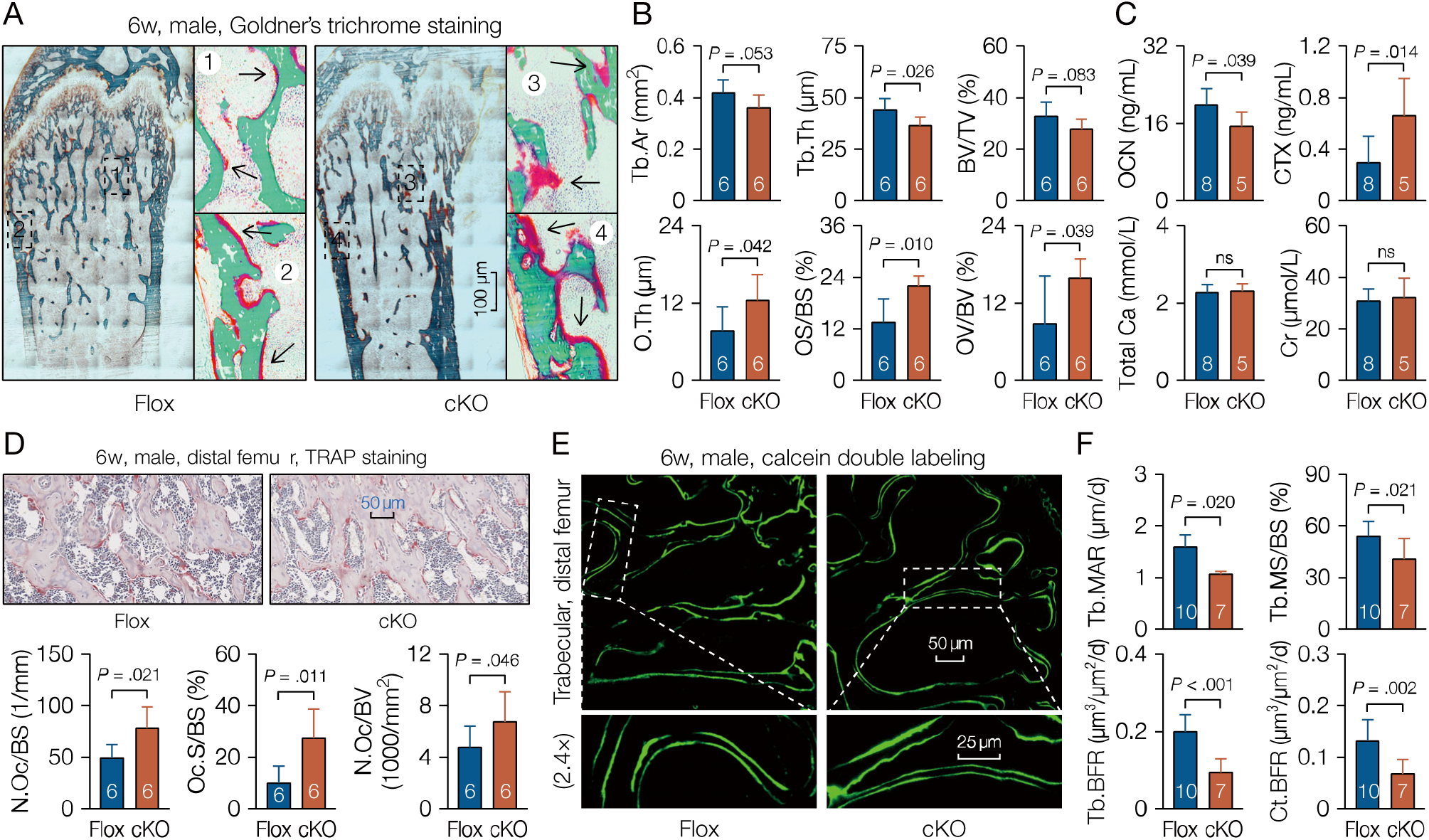
Low bone formation capacity in MACF1 cKO mice. (A) Representative images of Goldner’s trichrome staining showing mineralized matrix and osteoid in femur (male, 6-week old, coronal sections). Green shows mineralized bone, while red indicates osteoid. Numbered boxes are enlarged to show details. (B) Quantification of panel A. OV/BV, osteoid volume to trabecular bone volume. OS/BS, osteoid surface to trabecular bone surface. O.Th, average osteoid thickness. (C) Sandwich enzyme immunoassay for *in vitro* quantification of serological factors. OCN, osteocalcin; CTX, carboxy-terminal collagen crosslinks; Cr, creatinine. (D) Representative images of TRAP staining showing bone resorption in femur (male, 6-week old, coronal sections). N.Oc/BS, number of osteoclasts per bone surface; Oc.S/BS, osteoclast surface per bone surface; N.Oc/BV, number of osteoclasts per bone volume. (E) Representative images of calcein double labeling showing mineral apposition and bone formation in femur (male, 6-week old, coronal sections). Boxed regions are enlarged. (F) Quantification of panel E. MAR, mineral apposition rate; MS/BS, mineralizing surface per bone surface; BFR, bone formation rate per bone surface. Data are represented as mean ± s.d. Statistical significance were determined using student’s *t*-test.

Bone microenvironment changes with age, and accordingly, the regulation of MSCs osteogenic differentiation varies as well. To confirm age-related phenotypes of the MACF1 deficient mice, mice older than peak bone mass age (around 6 month) were examined. MicroCT analysis of 7-month old distal femur showed that in cKO mice, bone mass and bone volume were significantly lower than those in Flox mice (Figure 2F), together with deteriorated trabecular microarchitecture (Figure EV3B). Further, H&E staining of FFPE femur sections showed that in 12-month mice, trabecular bone volume of cKO mice was still lower (Figure 2G), and marrow adipocyte number increased in MACF1 deficient mice (Figure EV3C). It is worth noting that, with increased age, BV/TV disparity between Flox and cKO mice increased from 23.9% at 6w to 35.7% at 12m (Figure 2C, 2F and 2G), suggesting that loss of MACF1 impairs bone properties more seriously in aged mice. Together, these results suggest that MACF1 positively regulates bone mass and bone quality in mice, and that loss of MACF1 confers risk for osteoporosis and fracture.

### MACF1 is required for bone formation

Osteoblasts and osteoclasts are two main cell types in mature bone tissue that synthesize or decompose bone matrix to coordinate bone remodeling. To further understand the behavior of MACF1 in bone development, histochemistry approaches were utilized. Firstly, to differentiate mineralized bone from osteoid that undergoing rapid bone growth, undecalcified femur sections (plastic sections) from 6-week old male mice were prepared. The Goldner’s trichrome staining results showed that in addition to porotic and lesser trabeculae, the cKO mice also showed atypical accumulated osteoid on surfaces of both the trabecular and cortical bone (Figure 3A and 3B).

Certain substances produced during bone remodeling are secreted into peripheral blood, and could be used as biomarkers to reflect osteoblast function, osteoclast function and metabolic rate of the bone. We next quantified bone metabolism-related serological factors in same old male mice, and found that bone formation-related factors such as osteocalcin (OCN) were decreased in cKO mice, while bone resorption-related factors like carboxy-terminal collagen crosslinks (CTX) level were elevated (Figure 3C), suggesting reduced bone matrix mineralization and increased type I collagen decomposition due to MACF1 deletion. Furthermore, FFPE sections were also prepared and stained for tartrate-resistant acid phosphatase (TRAP). Consistent with earlier results, the cKO mice showed accumulated TRAP expression on bone surfaces (Figure 3D and S4A).

Lastly, to see the dynamic action of MACF1 on bone formation, mice at 6-week old and 3-month old were double-labeled with calcein, respectively. Fluorescence imaging of plastic sections showed that, regardless of mice age, femoral mineral apposition was slower in cKO mice (Figure 3E and S4B). Bone histomorphometric analysis further revealed that matrix mineralization and bone formation on bone surfaces were also slower (Figure 3F and S4B). These data suggest that MACF1 is required for bone formation in mice; insufficient MACF1 shifts the balance of bone remodeling and finally impairs bone formation.

### Loss of MACF1 disorganizes the cytoskeleton in MSCs

As a large structure protein, MACF1 interacts with microtubule and microfilaments simultaneously to coordinates cytoskeleton during cell proliferation and differentiation, and is reported to stabilize the cytoskeleton in different cell types (Fassett et al., 2013; Wang et al., 2017; Wu et al., 2008; Yue et al., 2016). To investigate the cytoskeletal role of MACF1 during osteogenic differentiation, MSCs from MACF1 deficient mice were isolated and analyzed. Immunofluorescence staining using antibody recognizing the alpha-tubulin showed in Flox cells distinct microtubule organizing center (MTOC), and ordered and well-organized microtubule filaments (Figure 4A), whereas in cKO cells, the MTs appeared to be acentric, MTOC disappeared, and the morphology of non-centrosomal microtubules became disorganized and cluttered (Figure 4A). Morphometric analysis of the microtubule demonstrated that mesenchymal deletion of MACF1 decreased anisotropy of MT filaments, and more importantly, depolarized orientation of the microtubule (Figure 5B), suggesting loss of MACF1 in MSCs impairs organization and structural regularity of the microtubule cytoskeleton. Next, we analyzed the actin cytoskeleton. Unlike the action of MACF1 to microtubule, loss of MACF1 did not seem to significantly affect the actin filaments (Figure 4C), similar anisotropy and orientation of the actin filament were found between cKO cells and Flox cells (Figure 4D).

**Fig. 4.**
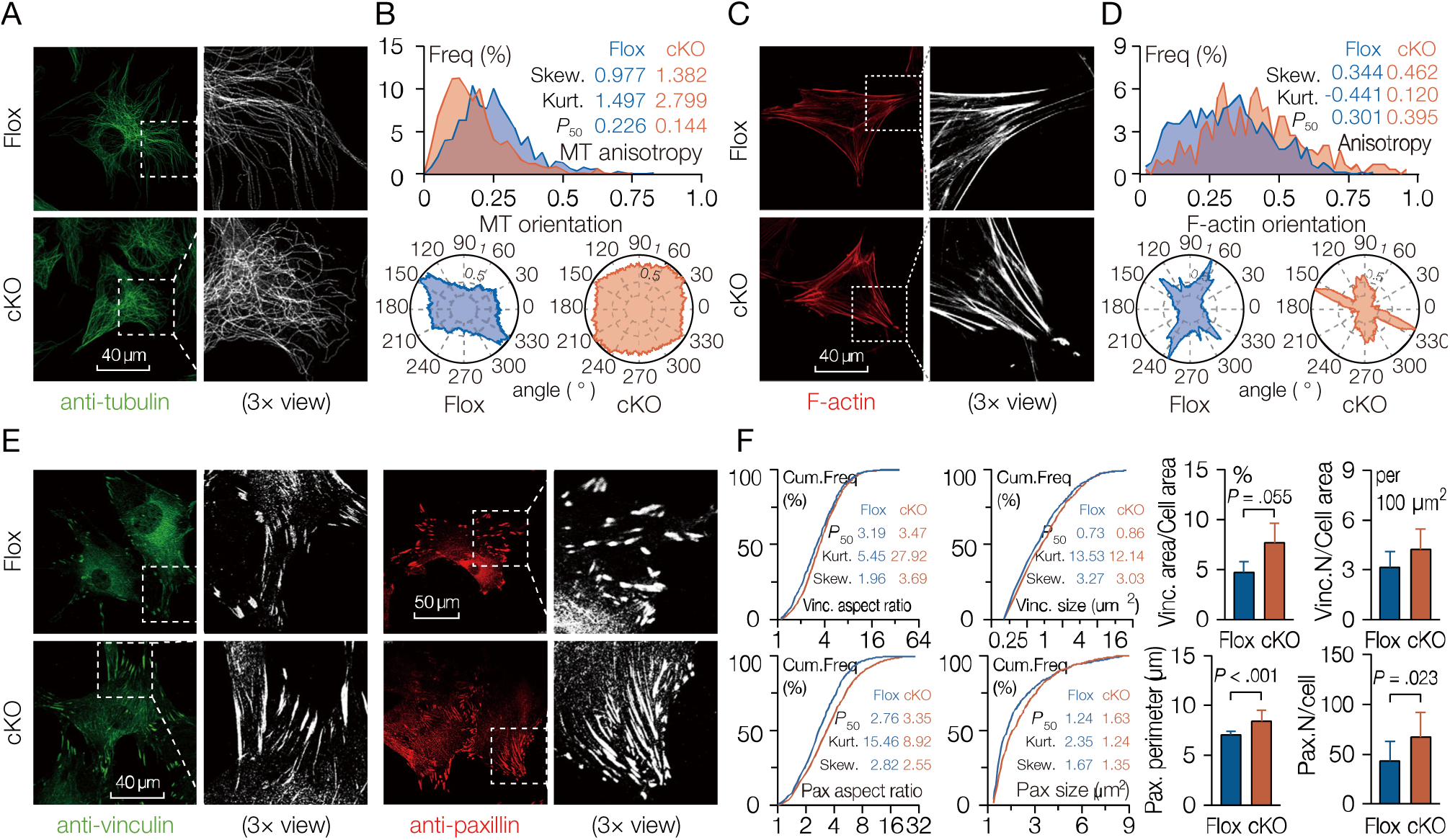
Loss of MACF1 alters the organization of cytoskeleton in MSCs. (A) Representative immunostaining images of MACF1 deficient MSCs stained with antibody recognizing alpha-tubulin. (B) Anisotropy and orientation of the microtubule cytoskeleton. MT, microtubule; Skew., skewness; Kurt., kurtosis; *P*_50_, median. (C) Representative immunostaining images of MSCs stained with rhodamine-conjugated phalloidin. (D) Anisotropy and orientation of the actin cytoskeleton. (E) Representative immunostaining images of MSCs stained with antibodies recognizing vinculin or paxillin to show focal adhesions. (F) Size and morphometric parameters of vinculin (Vinc.) or paxillin (Pax.) plaque. Cum.Freq, cumulative frequency. Data are represented as mean ± s.d. Statistical significance were determined using student’s *t*-test.

**Fig. 5.**
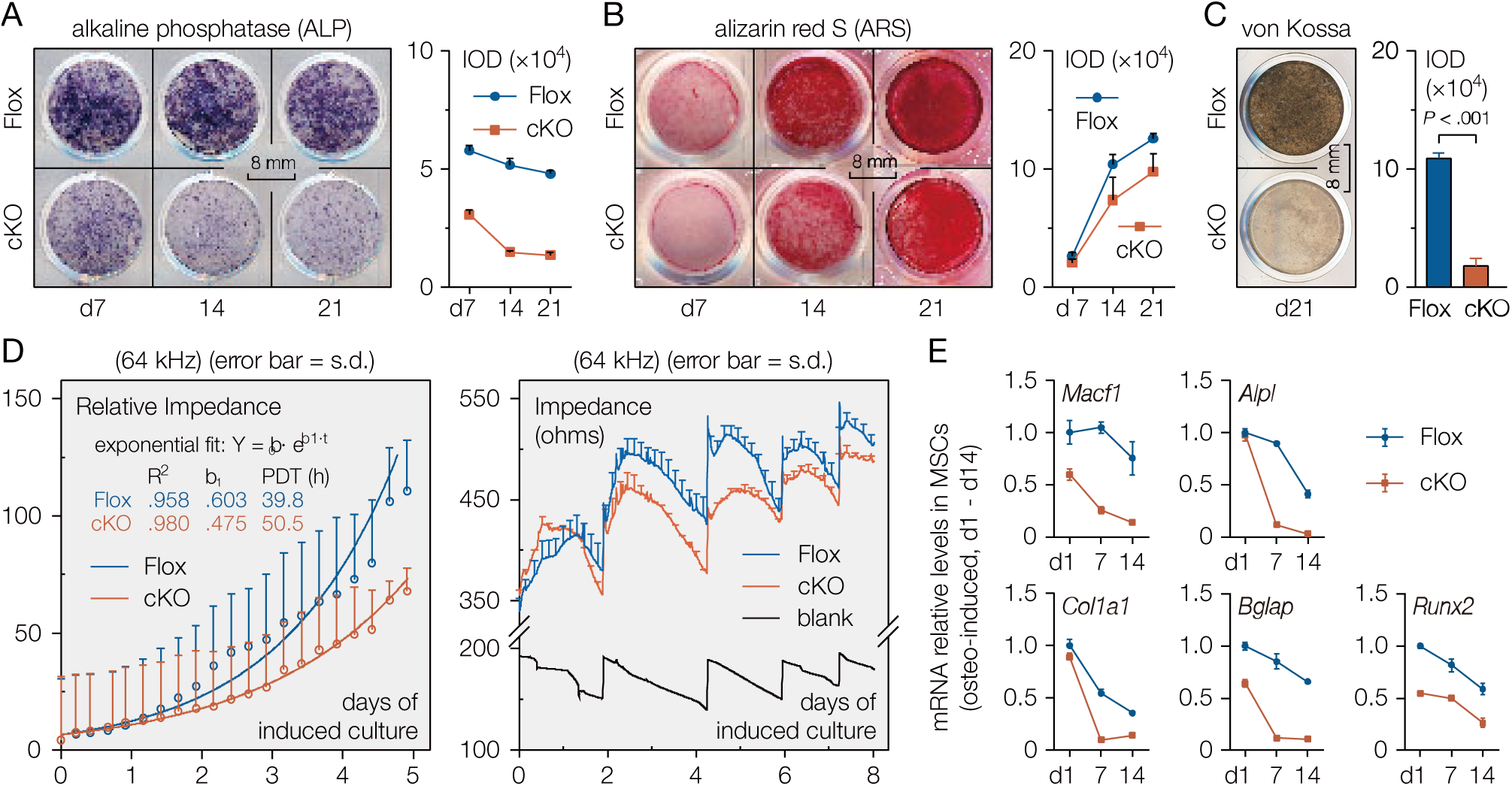
Loss of MACF1 inhibits osteoblastic differentiation in MSCs. (A) Representative images of von Kossa staining showing mineralization capacity after osteo-induced for 21 days. Staining intensity was quantified as IOD (integrated optical density). (B and C) Representative staining images showing alkaline phosphatase expression and mineralized nodules formation in MSCS during osteo-induced culture. (D) Real-time impedance curve showing proliferation and differentiation capability of MSCs during osteo-induced culture. PDT, population doubling time. (E) Real-time PCR analysis of osteogenic marker gene expression during osteo-induced culture. Data are represented as mean ± s.d.

Focal adhesions (FAs) are end parts of the F-actin that extend into the extracellular matrix. It is reported that MACF1 specifically promotes the crosslinking of microtubule and microfilament, and guide growth of the microtubule plus-ends towards FAs (Stehbens and Wittmann, 2012; Wu et al., 2008). To further understand the action of MACF1 on the cytoskeleton, FAs in MACF1 deficient MSCs were stained and analyzed. Immunofluorescence staining using antibodies recognizing vinculin or paxillin showed that, upon MACF1 deletion, the size of FAs increased significantly, and morphology of FAs changed from spindle-shaped in Flox cells to filament-like in cKO cells (Figure 4E and 4F), and the number of focal adhesions also increased in cKO cells (Figure 5F), indicating increased spread area of focal adhesions in cKO MSCs. These results suggest that MACF1 is necessary for maintaining microtubule integrity and focal adhesion morphology to coordinate cytoskeleton in MSCs.

### Loss of MACF1 inhibits osteogenic differentiation in MSCs

Although the *in vitro* roles of MACF1 have been studied in cells such as neuron and epidermis cell, much is still unknown about its *in vivo* functions in bone forming cells. To study the involvement of MACF1 in MSCs during osteogenic differentiation, we isolated and cultured bone-derived MSCs from the MACF1 cKO mice. Firstly, differentiation capability of MSCs during osteogenic induction was assayed. Von Kossa staining showed that at 21^st^ day of osteo-induced culture, loss of MACF1 greatly inhibited mineralization capability in MSCs (Figure 5A). In addition, alkaline phosphatase (ALP) and Alizarin Red S (ARS) staining further verified that during osteo-induction, MSCs lack of MACF1 had less ALP activity and less mineralized nodule formation (Figure 5B, C and S6A), indicating inhibited osteogenic differentiation capability.

Next, proliferation and differentiation capability of differentiating MSCs were detected non-invasively by electric cell-substrate impedance sensing (ECIS) assay. By monitoring real-time impedance values during osteo-induced culture, the ECIS curve showed that during exponential growth, proliferation capability decreased significantly in MACF1 deficient MSCs (Figure 5D, left), population doubling time increased from 39.8 h in Flox cells to 50.5 h in cKO cells. In addition, the ECIS curve reveled that loss of MACF1 significantly suppressed differentiation capability of MSCs, especially in early phase of osteogenic differentiation (Figure 5D, right).

Lastly, qPCR analysis revealed that, during osteogenic differentiation, levels of either *Macf1* or osteogenic genes decreased with induction time (Figure 5E), and more importantly, levels of both *Macf1* and osteogenic genes such as alkaline phosphatase (*Alpl*), collagenase 1 alpha 1 (*Col1a1*), osteocalcin (*Ocn*) and *Runx2* decreased significantly in MACF1 deficient cells (Figure 5E), suggesting impaired osteogenic differentiation capability upon MACF1 deletion. These data indicate that MACF1 is necessary for maintaining osteogenic differentiation, and lack of MACF1 would impairs osteogenic functions in MSCs.

### MACF1 interacts with SMAD7 in MSCs

To explore novel downstream molecules that interact with MACF1 to regulate osteogenic differentiation, we performed iTRAQ-labeled proteomics experiment using MC3T3-E1 preosteoblasts. First we co-immunoprepacited cell lysate using anti-MACF1 antibody, and confirmed immunoprecipitation capability of the anti-MACF1 antibody (Figure 6A, S7A and S7B). Next, the IP products were labeled with iTRAQ reagent and subjected to liquid chromatography followed by tandem mass spectrometry (LC-MS/MS) analysis. Unique peptide analysis showed that there were more than 60 proteins interacting with MACF1 in MC3T3-E1 cells, among which 5 were transcription factors (TFs), i.e., BHLHE40, HES1, REL, SMAD7, ZFP748 (Figure 6B). To further verify the involvement of these TFs in osteogenic differentiation, the ToppGene database was used to predict their potentials in osteogenic regulation, and highlighted SMAD7 as a most likely transcription factor in osteogenic regulation (data for ZFP748 not found) (Figure 6C, S7C and S7D). In addition, qPCR analysis revealed that in MC3T3-E1 cells, *Smad7* still had relatively high abundance, and in *Macf1* knockdown cells, *Smad7* levels was also lower (Figure 6D). Lastly, we isolated MSCs from wild type mice and performed co-IP to test SMAD7-MACF1 interaction in MSCs. The results showed that SMAD7 was detectable in the immunoprecipitated product, indicating interaction of MACF1 and SMAD7 in MSCs (Figure 6E). Besides, immunofluorescence showed from the perspective of intracellular distribution co-localization of MACF1 and SMAD7, which in turn confirmed the Co-IP result (Figure 6F).

**Fig. 6.**
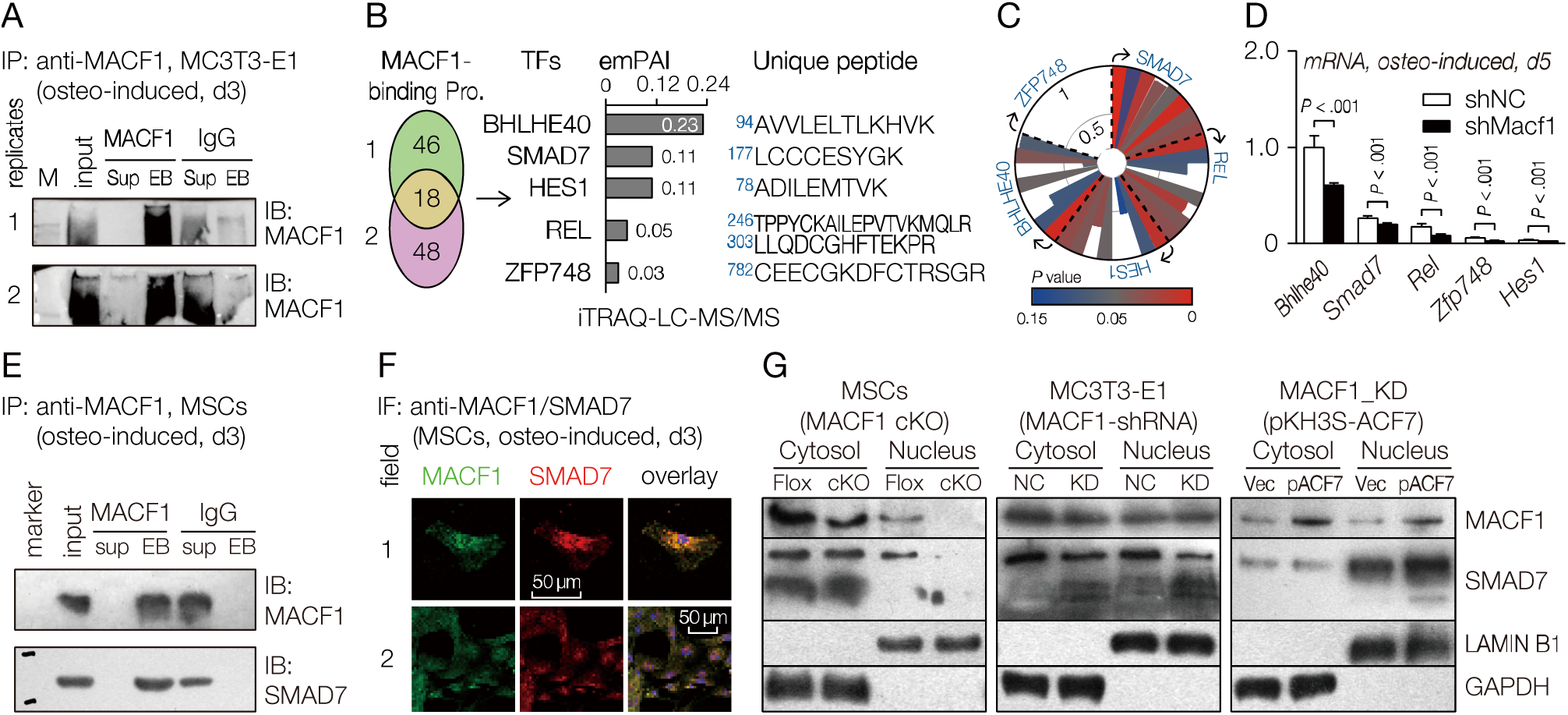
MACF1 interacts with SMAD7. (A) Co-immunoprecipitated products using anti-MACF1 antibody for iTRAQ analysis. (B) MACF1-interacting proteins identified by co-immunoprecipitation coupled LC-MS/MS assay, abundance of MACF1-interacting transcriptional factors (TFs) and unique peptide sequence (positions marked by Arabic numerals) were also provided. emPAI, exponentially modified protein abundance index. (C) Potential of candidate TFs to be involved in osteogenic differentiation. Radial coordinates represent the *Score* values predicted by ToppGene database, while angular coordinates show 8 predicted items (clockwise: GO: Molecular Function; GO: Biological Process; GO: Cellular Component; Human Phenotype; Mouse Phenotype; Pathway; Pubmed; Disease). (D) Real-time PCR analysis of candidate TFs in MACF1 knockdown MC3T3-E1 preosteoblasts after osteo-induced for 5 days. (E) Co-immunoprecipitation assay to detect the association of MACF1 with SMAD7. MSCs were co-immunoprecipitated using anti-MACF1 antibody or IgG. (F) Expression and distribution of MACF1 and SMAD7 in normal MSCs after induced. (G) Western blot analysis of MACF1 and SMAD7 levels in MACF1 cKO MSCs, MACF1 knockdown preosteoblasts, and MACF1 stable transfected MACF1 knockdown preosteoblasts. ACF7 (Actin cross-linking family protein 7) is a synonym for MACF1. Lamin B1 and GAPDH were used as internal reference for nucleus and cytoplasm, respectively. Data are represented as mean ± s.d. Statistical significance were determined using student’s *t*-test.

On the basis of MACF1-SMAD7 interaction, to further clarify their regulatory pattern in osteogenic differentiation, we manipulated MACF1 expression in a variety of cell lines and tested SMAD7 expressions in both the cytoplasm and nucleus. Firstly, in Floxed MSCs, we found that MACF1 was mostly expressed in the cytoplasm, and upon MACF1 deletion, expression of MACF1 decreased significantly in both the cytoplasm and nucleus. Similar to the MACF1 expression pattern, SMAD7 also expressed mostly in cytoplasm, and deletion of MACF1 significantly decreased expression of SMAD7, especially in the nucleus (Figure 6G, left). Next, we constructed a *Macf1* knockdown (MACF1-shRNA) MC3T3-E1 cell line, and found in KD cells reduced expression of SMAD7 (Figure 6G, middle). Lastly, we constructed a cell line to restore MACF1 expression by transfection of pKH3S-ACF7 plasmid into the *Macf1* knockdown cells, and found that, as compared with empty plasmid control group, restoration of MACF1 significantly elevated SMAD7 levels, especially in the nucleus (Figure 6G, right). These data suggest that, in bone forming cells, SMAD7 interacts with MACF1 and shares a similar expression trend.

### MACF1 facilitates nucleus translocation of SMAD7 to drive osteogenic differentiation

SMAD7 is a key member of the TGFβ superfamily SMAD family that regulates cell metabolism. Studies show that, SMAD7 may play versatile roles in cell signaling. Cytoplasmic SMAD7 was found to inhibit cellular functions (e.g. in bone forming cells) (Wu et al., 2016), while nuclear SMAD7 was reported to promote cellular functions (e.g. in myoblasts) (Miyake et al., 2010). Still, there’s no data to clarify the nuclear roles of SMAD7 in bone forming cells. To further clarify how SMAD7 participates in osteogenic regulation through interacting with MACF1, we constructed a conventional SMAD7 overexpression plasmid (pcDNA_SMAD7) and a nuclear localization signal-carrying SMAD7 overexpression plasmid (pcDNA_SMAD7-NLS), respectively, to studied the subcellular functions of SMAD7 in MC3T3-E1 osteoblasts (Figure 7A).

**Fig. 7.**
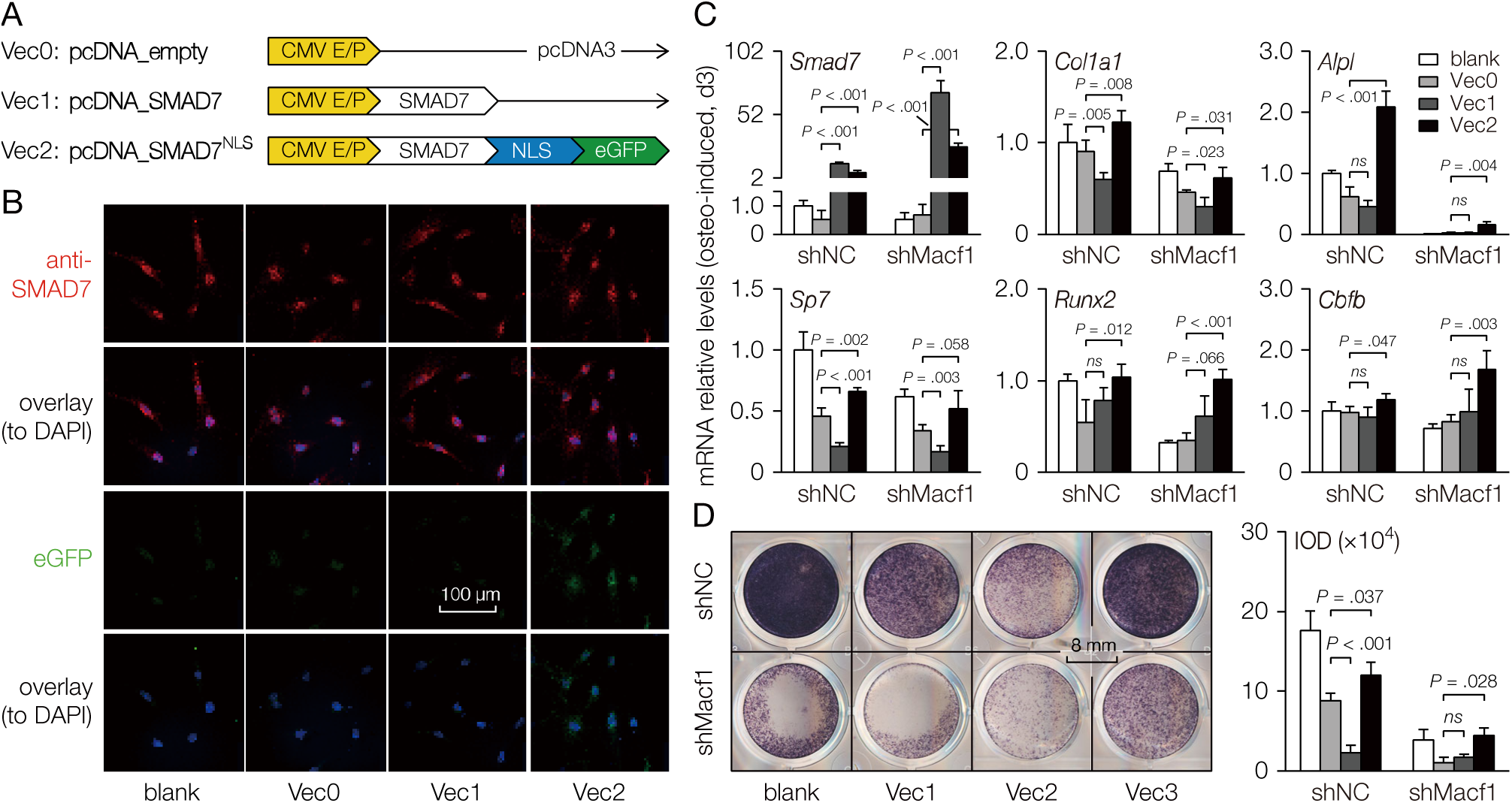
MACF1 facilitate SMAD7 nucleus translocation to drive osteoblastic differentiation. (A). Schematic diagram of the SMAD7 expression plasmids. The mouse SMAD7 coding sequence was cloned into pdDNA3 backbone. NLS, nuclear localization sequence (SV40 large T antigen). (B) Representative immunostaining images of MC3T3-E1 preosteoblasts showing SMAD7 (red) and eGFP (green) expression after plasmid transfection. Blank group was not transfected. (C) Real-time PCR analysis of relative gene expression of Smad7, osteogenic marker genes and osteogenic-related TFs in *Macf1* knockdown MC3T3-E1 preosteoblasts after plasmid transfection. (D) Representative staining images showing alkaline phosphatase expression in *Macf1* knockdown MC3T3-E1 preosteoblasts after plasmid transfection. Data are represented as mean ± s.d. Statistical significance were determined using student’s *t*-test.

Next, the transfection efficiency was examined. Immunofluorescence images showed that, 12 h after transfection, level of cellular SMAD7 increased, in cells transfected with conventional plasmid, SMAD7 distributed mostly within the cytoplasm, while in cells transfected with NLS-carrying plasmid, SMAD7 concentrated mostly within the nucleus (Figure 7B). Then, we explored the effects of SMAD7 overexpression on osteogenic differentiation. qPCR analysis revealed that, as compared with empty plasmid transfected group (Vec0), *Smad7* expression increased dramatically after transfected with two kinds of plasmids. In cells transfected with conventional SMAD7 plasmid, expressions of osteogenic genes such as *Col1a1* and *Alpl* decreased simultaneously, levels of osteogenic-related transcription factors reduced as well (Figure 7C). However, in cells transfected with NLS-carrying SMAD7 plasmid, levels of both osteogenic genes and transcription factors increased significantly (Figure 7C and S8A).

Lastly, cells transfected with plasmids were osteo-induced and stained for ALP. The results showed that, as compared with the empty plasmid transfected group, ALP activity decreased in conventional SMAD7 plasmid transfected cells, and increased in NLS-carrying plasmid transfected cells (Figure 7D). These results suggest that, during the process of osteogenic differentiation, cytoplasmic SMAD7 plays an inhibitory role, while nuclear SMAD7 positively regulates osteogenic functions. Concerning the evidence that SMAD7 interacts with MACF1, these data indicate that during osteogenic differentiation, MACF1 facilitates nuclear translocation of SMAD7 to drive downstream osteo-related signaling and to positively regulate bone formation.

## DISCUSSION

In this study, we identified MACF1 as a key regulator that controls osteogenic functions in mice. We show that MACF1 is necessary for bone development and formation. Loss of MACF1 decreases bone mass and quality in adult mice, which confers risk for degenerative bone diseases such as fracture. In addition, MACF1 stabilizes the microtubule cytoskeleton and controls focal adhesion morphology and regulates osteogenic differentiation in MSCs. Although the role of MACF1 in material transportation has been reported, a novel transcriptional factor SMAD7 that interacts directly with MACF1 is identified until this study, which initiates and promotes downstream osteogenic pathways. Our findings provide a new mechanism for understanding osteogenic differentiation and bone formation regulation, and identified MACF1 as a potential therapeutic target for degenarative bone diseases such as osteoporosis.

### Essential roles of MACF1 in regulating bone development and formation

Conventional or nervous-tissue-specific knockout of MACF1 was reported to be embryonic lethal (Chen et al., 2006; Goryunov et al., 2010), but mesenchymal-specific MACF1 KO mice survived and are physically normal, suggesting a relatively independent role of MACF1 in bones. However, mice lacking MACF1 show different degrees of retardation in skull and long bone development. Considering the MACF1 deficiency phenotypes found in other tissues (Fassett et al., 2013; Goryunov et al., 2010; Ka et al., 2014; Ka and Kim, 2016; Liang et al., 2013; May-Simera et al., 2016), we believe that MACF1 is also responsible for positive regulation of bone growth and development. Loss of MACF1 would likely cause abnormalities during early stages of bone development.

In adult mice, especially with increased age, loss of MACF1 impairs bone mass, mineral density and bone properties tremendously, which experimentally confirmed MACF1 as a potential key regulator for degenerative bone disorders such as osteoporosis. Further investigations show that, loss of MACF1 inhibits bone surface mineralization capability, promotes osteoclasts functions and subsequently inhibits bone formation. Therefore, we propose that loss of MACF1 disrupts the balance of bone remodeling, and confers risk for degenerative bone diseases. Interestingly, MACF1 deficient mice have more osteoid deposited on bone surfaces, which is inconsistent with what we have hypothesized. Further serological analysis shows that MACF1 deficient mice have less ability to mineralize osteoid, elevated osteoid is likely to be produced in a compensatory manner, but insufficient MACF1 leads to insufficient mineralization capability, and finally leads to reduced bone mass.

### Loss of MACF1 impairs cytoskeleton dynamics and osteogenic functions

As an important cytoskeleton regulator (Suozzi et al., 2012), we hypothesized that MACF1 also regulates the cytoskeleton in MSCs. Our results show that, loss of MACF1 disorganized the microtubule cytoskeleton and depolarized the cell. Previously, it is reported that loss of MACF1 disturbs the microtubule cytoskeleton in epidermis, neuron and myoblasts, therefore we believe MACF1 is also important for maintaining the microtubule cytoskeleton in MSCs. The actin cytoskeleton is not the same, however, loss of MACF1 doesn’t seem to affect the morphology or polarity of the actin filaments. Similar phenomenon was found in epithelial cells, in which MACF1 deletion affected the microtubule cytoskeleton, rather than the microfilaments, and it is the microfimants, rather than the MTs, that could reorganize themselves in the absence of MACF1 (Kodama et al., 2003). Considering the molecular structure of MACF1, a tentative inference on this difference is that MACF1 crosslinks differently with microtubules and microfilaments, respectively. Focal adhesions are essential parts of the cytoskeleton, we find that MACF1 deficient MSCs have larger and more focal adhesions. Earlier studies also show that loss of MACF1 stabilizes focal adhesions, impairs its dynamic characteristics and then affects cell motility (Ning et al., 2016; Wu et al., 2008; Yue et al., 2016). Taking together these results, we propose that MACF1 regulates focal adhesion morphology to control MSCs adhesion and motility.

### Interaction of MACF1 and SMAD7 during osteogenic differentiation

MSCs differentiate into bone forming cells to support osteogenic functions. Based on the *in vivo* results, we hypothesized that loss of MACF1 impairs osteogenic functions in MSCs. The *in vitro* experiments validate that osteogenic differentiation capability in MSCs rely heavily on sufficient expression of the *Macf1* gene. There are also more studies to show that loss of MACF1 inhibits cell differentiation (Ka and Kim, 2016), which are attributed to disorganized cytoskeleton (Ka et al., 2014). Next, we identified SMAD7 as a downstream transcriptional factor that interacts with MACF1 to be translocated into the nucleus to initiate downstream osteogenic pathways. SMAD7 has been shown to regulate osteogenic functions. A prevailing view is that SMAD7 act as a repressor in the TGF-β/BMP signaling pathway (Miyazawa and Miyazono, 2017; Wu et al., 2016), but emerging evidences are presenting its positive roles in osteogenic regulation (Li et al., 2014). We noticed that this may be due to its subcellular localization, but the nuclear roles of SMAD7 are still unknown. On the one hand, SMAD7 exists simultaneously in the cytoplasm and nucleus in different types of cells. For instance, nuclear SMAD7 in myoblasts promotes myogenesis (Miyake et al., 2010), similar results are also found in fibroblasts and other cells (Emori et al., 2012). On the other hand, MACF1 is reported to have ATPase activity (Wu et al., 2008), and mediate intracellular material trafficking (Chen et al., 2006; Slep et al., 2005), our data show that SMAD7 has multiple phosphorylation sites (Hornbeck et al., 2015), and MACF1 shuttles between the cytoplasm and nucleus. For this reason, we overexpressed SMAD7 in MSCs, and found nuclear-specific overexpression of SMAD7 promotes osteogenic differentiation, while whole cell overexpression (in fact mostly accumulated in the cytoplasm) shows inhibitory effects. By differentiate its subcellular roles, we uncover new function of SMAD7 in osteogenic differentiation regulation.

In summary, we discovered new roles of MACF1 in regulating osteogenic differentiation and bone formation, loss of MACF1 in bone forming cells impairs osteogenic functions and then leads to degenerative bone disorders such as osteoporosis. Further, we identified SMAD7 as a key transcriptional factor that interact with MACF1 to be translocated into the nucleus to initiate downstream osteogenic pathways.

## MATERIALS AND METHODS

### Generation of the MACF1 cKO mouse

Mice carrying the floxed MACF1 alleles (in B6 background) were acquired through a material transfer agreement between the University of Chicago (USA) and the Northwestern Polytechnical University (China). Construction of the targeting vector was reported previously (Wu et al., 2008), in which the loxP sites were integrated to excise exon 11, 12 and 13 of the *Macf1* gene (while in the original literature, the flanked exons were reported as E6 and E7) (Figure EV1A). Mice expressing Cre recombinase under the control of Prx1 promoter (in B6 background) were acquired from Biocytogen Co., Ltd. (Beijing, China). The MACF1 conditional knockout mice were generated by breeding the MACF1 Flox mice to the Prx1-Cre mice. Offspring genotype was determined by PCR analysis of genomic DNA from the toe tissue (or for embryos, from liver tissue).

Feed and water used for mice husbandry were pre-processed properly (^60^Co irradiated feed; boiled water), and were all available *ad libitum* to mice. Breeding and number of mice were controlled according to the 3Rs principle. Mice used for planned experiments were euthanatized after overdose injection of pentobarbital sodium (i.p., 1 g/Kg), while others were euthanatized by CO_2_ inhalation. All mice experiments were performed according to the *Guide for the Care and Use of Laboratory Animals* and under the permission of Laboratory Animal Ethics & Welfare Committee of the Northwestern Polytechnical University.

### Study design & animal grouping

To study the *in vivo* functions of MACF1, age-matched same-sex controls were set up (mice numbers are indicated in each figure). Unless otherwise stated, the MACF1^f/f^;Prx1^Cre/+^ mice (cKO) were considered to have the *Macf1* gene deleted specifically in MSCs, while the MACF1^f/f^ mice (Flox) were used as controls. All mice were maintained in barrier condition (SPF lab, 12/12 h light cycle, 24±1 °C, 30-70% RH) with no more than 4 in each cage.

### Cell preparation

Mouse mesenchymal stem cells were isolated from compact bone of 6-week old male mice (Zhu et al., 2010). Briefly, mice femora were isolated, cleaned and transferred to a 60-mm culture dish containing 5 mL α-MEM (supplemented with 0.1% penicillin/streptomycin and 2% FBS). Epiphyses in both ends were cut open for complete removal of bone marrow. Bone shafts were then excised into chippings and digested (α-MEM + 10% FBS + 1 mg/mL Type II collagen, 37 °C, 200 rpm, 1h), bone chips were then collected and seeded in a 60-mm dish for culture (37 °C, 5% CO_2_). Culture medium was changed on the third day to remove loosely attached cells and tissue debris. Cells at passage 2 to 5 were used for experiments.

Construction of the MACF1 knockdown MC3T3-E1 cell line (KD), and transfection of pKH3S-ACF plasmid into MACF1 KD cells were reported previously (Hu et al., 2015; Zhang et al., 2018).

Growth medium (10% FBS + 1% pen-strep + 1% L-glutamine in α-MEM) were used for subculture, and osteo-induction medium (10% FBS + 1% pen-strep + 1% L-glutamine + 0.1 μM dexamethasone + 50 μg/mL ascorbic acid + 10mM glycerol in α-MEM) were used for osteo-induction.

### RNA and qPCR

High quality total RNA from cell and tissue samples were isolated using TRIzol™ reagent according to the manufacturer’s instructions. RNA quality was determined by ultraviolet spectrophotometry. cDNA was synthesized on 1 μg total RNA using the cDNA synthesis kit (TaKaRa, Japan) following the manufacturer’s instructions. Real-time quantitative PCR was performed using the SYBR^®^ Green method on 2 μL cDNA (1:50) in a 20 μL PCR system. PCR data were analysis with the comparative CT method (2^−ΔΔ*C*^T). *Gapdh* or 18s was used as internal controls.

### Protein and western blot

For extraction of protein from bones, femora were excised free of connecting tissue, flash-frozen and pulverized into bone chips using a pestle and mortar in the presence of liquid nitrogen (while soft tissues were pulverized using a tissue pulverizer (PRIMA, PB100-SP08)), and then digested with protein lysis buffer (Beyotime P0013, supplemented with 1× protease inhibitor cocktail). The lysate was then centrifuged at 12 000 g, 4 °C for 2 minutes and the supernatant was collected. For fractionation of cytoplasmic and nuclear proteins, cell cultures were washed and lysed using the Nuclear and Cytoplasmic Protein Extraction Kit (Beyotime P0028) following the manufacturer’s instructions. Protein concentrations were determined using the BCA method.

For western blot analysis, protein samples separated by SDS-PAGE were transferred onto PVDF membranes and probed with antibodies against target proteins. Briefly, 20 μg protein samples were electrophoresed at 100V on 5% stacking gel/10% separating gel. After transferred to PVDF membrane (100V, 90 min), and incubated with 5% non-fat milk for 30min, the membrane was probed with primary antibodies against MACF1 (1:1000), SMAD7 (1:1500), Lamin B1 (1:2000), and GAPDH (1:2000). Washed membranes were then incubated with HRP-conjugated secondary antibodies, and developed and imaged (Gel Doc^TM^ XR+ Gel System).

### Immunofluorescence & Cytoskeleton staining

Immunofluorescence staining was performed on fibronectin-coated coverslips. Briefly, Cells were plated on coverslips (1×10^4^/cm^2^) and cultured under normal condition for 12h. Then cells were fixed with 4% PFA and rinsed 5 min with TBS, 10 min with TBS-0.5% TritonX-100 (TX), and 3×5 min with TBS-0.1%TX, and then blocked with 2% BSA for 10 min. Primary antibodies were added onto each coverslip (10 μL, 1:50), and incubated at 4 °C overnight. The next day, after washed with TBS-0.1%TX, fluorescein-labeled secondary antibodies were added (10 μL, 1:100), and incubated for 1 h. Nucleuses were stained using 5μg/ml DAPI. F-actin were stained using rhodamine phalloidin (ThermoFisher, R415). Mounted coverslips were then examined by confocal microscopy (Leica TCS SP8, Germen). Fluorescent images were analyzed with the FibrilTool (Boudaoud et al., 2014), CytoSpectre (Kartasalo et al., 2015) and Image-Pro Plus.

### Co-immunoprecipitation and iTRAQ-LC-MS/MS

Co-immunoprecipitation was performed using freshly-harvested MC3T3-E1 cells incubated with antibody recognizing the MACF1 protein. Briefly, cells were harvested in RIPA buffer. Whole cell lysates were incubated at 4°C overnight with 4 µg anti-MACF1 antibody (Abcam, ab117418) or 2 µg Rabbit IgG (Abcam, ab46540). The antigen/antibody complexes were then mixed with 40 µL recombinant Protein A+G agarose beads (Beyotime, P2055), and incubated for 2 h at room temperature. Immunocomplexes were then centrifuged at 1000 g for 5 min to remove supernatant, and then beads complexes were washed with RIPA buffer for 5 times. After washing, beads complexes were resuspended with 20 µL 1X SDS-PAGE electrophoresis loading buffer, and bound proteins were retrieved from agarose beads by boiling for 5 min.

For analysis of MACF1-interacting proteins, co-immunoprecipitated products were labeled with iTRAQ reagents and subjected to LC-MS/MS analysis.

### Cleared skeleton preparation

Fetuses or neonatal mice were euthanized by anesthetization (pentobarbital sodium, i.p., 0.5 g/Kg). Samples were skinned and eviscerated before being fixed in 95% ethanol for 24 h, and then incubated in acetone overnight. To stain for cartilage, samples were rinsed with water for 3 changes and left in 0.03% (w/v) Alcian blue solution for about 18 h (depending on the development stage of the samples), and then cleared in 1% KOH until soft tissues were hardly visible. To visualize calcified bone, samples were counterstained in 0.02% Alizarin Red S solution for about 12 h, and then cleared in 1% KOH. Images were acquired by a digital camera, and dimensional parameters were acquired using Image-Pro Plus.

### Bone Densitometry & Micro-computed tomography

Bone mineral density was measured using a high resolution dual-energy X-ray imaging system (MEDIKORS, InAlyzer, 55KeV and 80Kev). Microarchitecture and stereological parameters were acquired using an micro-computed tomography instrument (EL-SP, GE, 80 kPV, 80 μA, 3000 ms) at an isotropic resolution of 8 μm. Data were then reconstructed at a voxel size of 16×16×16 μm. Stereological parameters of the trabeculae were measured within a 1 mm-high region above the distal femur epiphyseal plate using MicroView (a GE distribution). Sample scanning and data analysis adhered to a previous protocol (Bouxsein et al., 2010). Additional parameters were acquired using ImageJ and Image-Pro Plus.

### Three-point bending test

Mechanical properties of the femur were tested by means of 3-point bending using an universal test machine (Instron 5943). Briefly, freshly-dissected femora were cleaned free of connective tissue and wrapped in normal saline-soaked gauze and kept moist at 4 °C. For subsequent test, femora were secured to the lower supporting pins (span: 9 mm, fillet radius: 1 mm) one by one. After preloaded with 1N for 5 s, the tissues were loaded to failure at 1.5 mm/min, and load-displacement curves were collected. The inner and outer diameters of the loaded femora were measured using a 3D digital microscope (Hirox KH-8700, Japan). Mechanical parameters were acquired using a built-in software and a previous reported method (Sharir et al., 2008).

### Histochemical, calcein labeling and histomorphometric analysis

Formalin-fixed paraffin-embedded (FFPE) tissue sections were prepared and stained using conventional methods. Briefly, fresh tissue were fixed in 4% PFA, decalcified in 10% EDTA (for femoral FFPE sections only), dehydrated through graded ethanol, and embedded into paraffin. Sections were obtained at 4 μm. HE, ALP staining (0.5% β-glycerophosphate Disodium + 0.5% Barbital sodium + 0.5% CaCl_2_ + 0.5% Mg_2_SO_4_ + 2% cobalt nitrate + 1% ammonium sulfide) and TRAP staining (50mM C_4_H_12_KNaO_10_) were performed on serial sections. Stained sections were then scanned using a digital slide scanner (Aperio AT2, Leica, German).

For plastic embedded sections, 6-week-old mice were injected intraperitoneally with calcein (10 mg/kg) 10 days and 2 days prior to euthanasia, respectively. Femora were then collected and fixed, and embedded in gradient methyl methacrylate after conventional tissue dehydration and clearing. Sections (∼20 μm) were prepared using the EXAKT 300CP tissue cutting systems and stained using modified Goldner’s trichrome staining method (Gruber, 1992). Slides were visualized using a phase contrast microscope (OSTEO-SCAN). For histomorphometric parameters, images were analyzed using Histomorph suite (van’t Hof et al., 2017) and Image-Pro Plus.

### Serological factors

Whole blood collected by cardiac puncture was used for serum separation. Serological factors were detected by means of enzyme-linked immunosorbent assay following the manufacturer’s instructions (Cloud-Clone Corp.). O.D. values were read at 450 nm (BioTek Synergy HT, VT, USA).

### Cell culture staining

Von Kossa staining was performed using 5% AgNO_3_ after being induced for 21 days. ALP staining was performed using a BCIP/NBT alkaline phosphatase color development kit (Beyotime, C3206) following to the manufacturer’s instructions. Briefly, cell cultures were washed with cold PBS, fully fixed in 4% PFA and washed. The staining working solution was prepared by mixing BCIP (5-Bromo-4-chloro-3-indolyl phosphate, 300×) and NBT (nitro blue tetrazolium chloride, 150×) into the color developing buffer, cell cultures were then stained in the dark for 15 min. Nodules formation staining was performed using 0.5% Alizarin Red S solution (ARS). Stained cell culture plates were scanned using a digital scanner (CanoScan 9000F MarkII, Japan).

### Electric Cell-Substrate Impedance Sensing

Dynamic physiological properties of MSCs during proliferation and differentiation were monitored non-invasively using the ECIS Zθ system (Applied BioPhysics, Troy, NY) (Bagnaninchi and Drummond, 2011). Briefly, cells plated onto fibronectin-coated (20 μg/mL) standard 8-well array (8W10E+) were supplemented with 400 μL growth medium, and cultured in CO_2_ incubator for 6 h before initial test. To measure impedance values of differentiating MSCs, growth medium was replaced with osteogenic osteo-induction medium, and data were collected every 80 s at a current frequency of 64 kHz.

### Plasmid preparation

The SMAD7 overexpression vector was cloned into the pcDNA3 backbone. Briefly, double enzyme digestion was performed on selected restriction enzyme sites. Then, seamless clone-specific primers were designed to match selected restriction enzyme sites, and the SMAD7 CDS sequence was amplified by PCR. Products from the aboved steps were purified by gel extraction. Linearized vector and CDS insert were recombined in the presence of high-fidelity polymerase. The recombinants were then transformed into Stbl3 competent cells and sequenced. Endotoxin-free plasmid was prepared and transfected into cells using nano particle transfection reagent (Engreen H4000) according to the manufacturer’s instructions.

### Data processing & Statistics

Image acquisition equipments include phase contrast microscope, fluorescence microscope, confocal microscope, digital camera, digital scanner, digital slide scanner, gel imaging system, micro-computed tomography, and dual-energy X-ray absorptiometry. To ease observation and statistical analysis, certain images may have been pre-processed using Adobe^®^ Photoshop^®^. Operations applied to image data include cropping, scaling, flipping, background clearing, local zoom in and tagging. Other lossy operations such as brightness adjustment, contrast adjustment, levels adjustment were applied only to a limited number of images simultaneously, and great efforts were made to ensure that these operations did not change the essence or any measurable variables of the images.

All data are presented as mean ± s.d. Statistical significance between two groups were determined using Student’s *t*-test. A *P*-value less than 0.05 was considered statistically significant.

## ACKNOWLEDGMENTS

The authors would like to thank Dr. Hong Zhou (the University of Sydney), Dr. Changqi C. Zhu (Ferris State University), Dr. Yi-Ping Li (the University of Alabama at Birmingham) and Dr. Li Sun (Icahn School of Medicine at Mount Sinai) for technical guidance. This work was supported by National Natural Science Foundation of China (31400725, 31570940, 81772017), Shenzhen Science and Technology Plan (JCYJ20160229174320053), grant BKJ17J004, China Postdoctoral Science Foundation (2017M610653), and Basic Research Foundation of NPU (3102016ZY037).

## AUTHOR CONTRIBUTIONS

Conceptualization, F.Z., X.M. and A.Q.; Methodology, F.Z., X.M., W.Q., Z.C., P.H., C.Y.; Formal Analysis, F.Z., J.M.; Investigation, F.Z., X.M., W.Q., P.W., R.Z., Z.C., P.H., L.C., C.Y.; Writing-Original Draft, F.Z.; Supervision, A.Q., G.Z.; Project Administration, A.Q., G.Z., Y.L.; Funding Acquisition, A.Q., L.H., Y.T., X.L.

## Working model

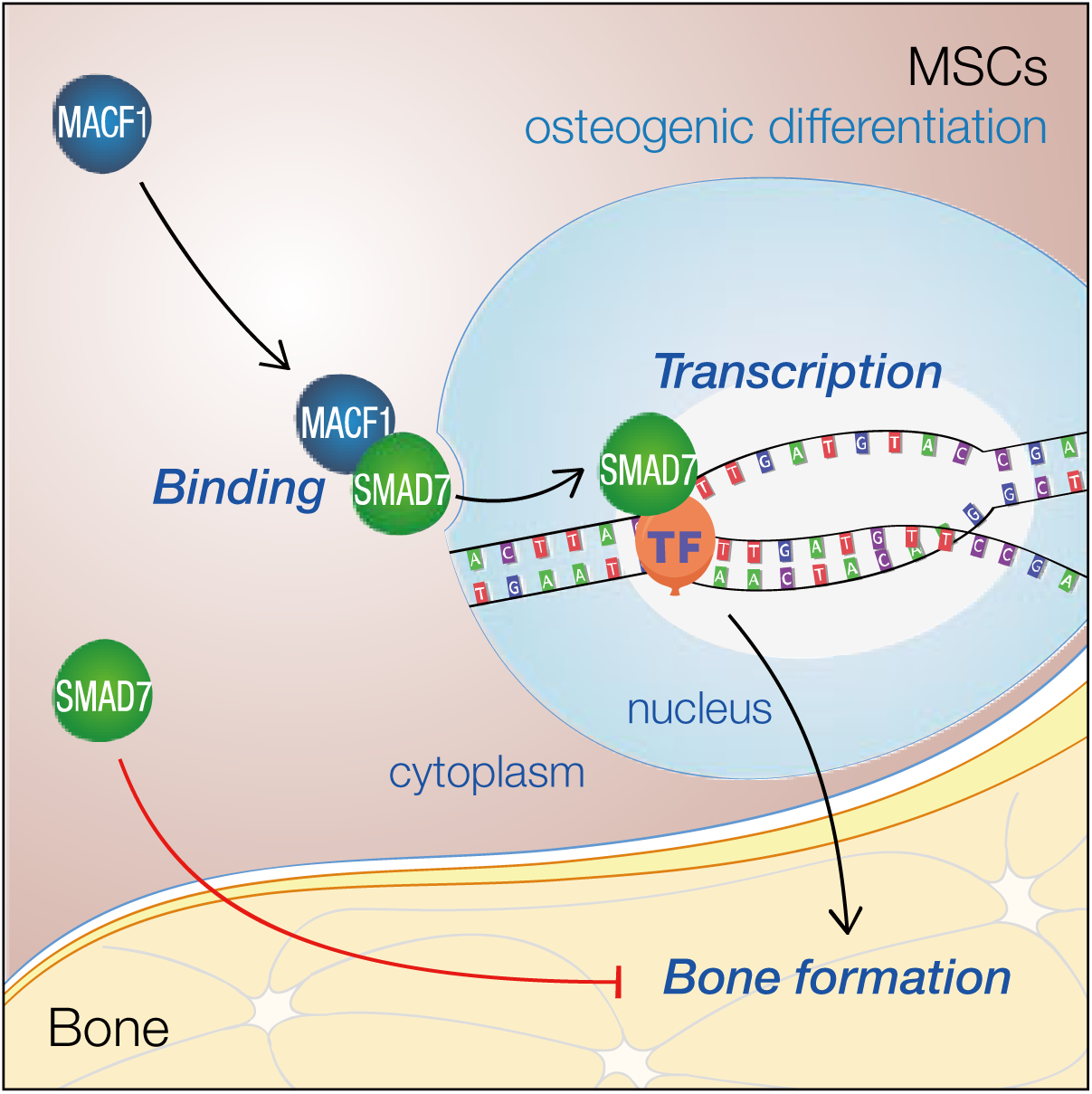
Working model. MACF1 is essential for maintaining osteogenic differentiation and bone formation. In MSCs, cytoplasmic SMAD7 inhibits osteogenic differentiation, while MACF1 promotes nuclear translocation of SMAD7 to initiate osteogenic pathways.

