## Supplemental Figures for "MACF1 Facilitates SMAD7 Nuclear Translocation to Drive Bone Formation in Mice"

### SUPPLEMENTARY FIGURE LEGENDS

Figure S1

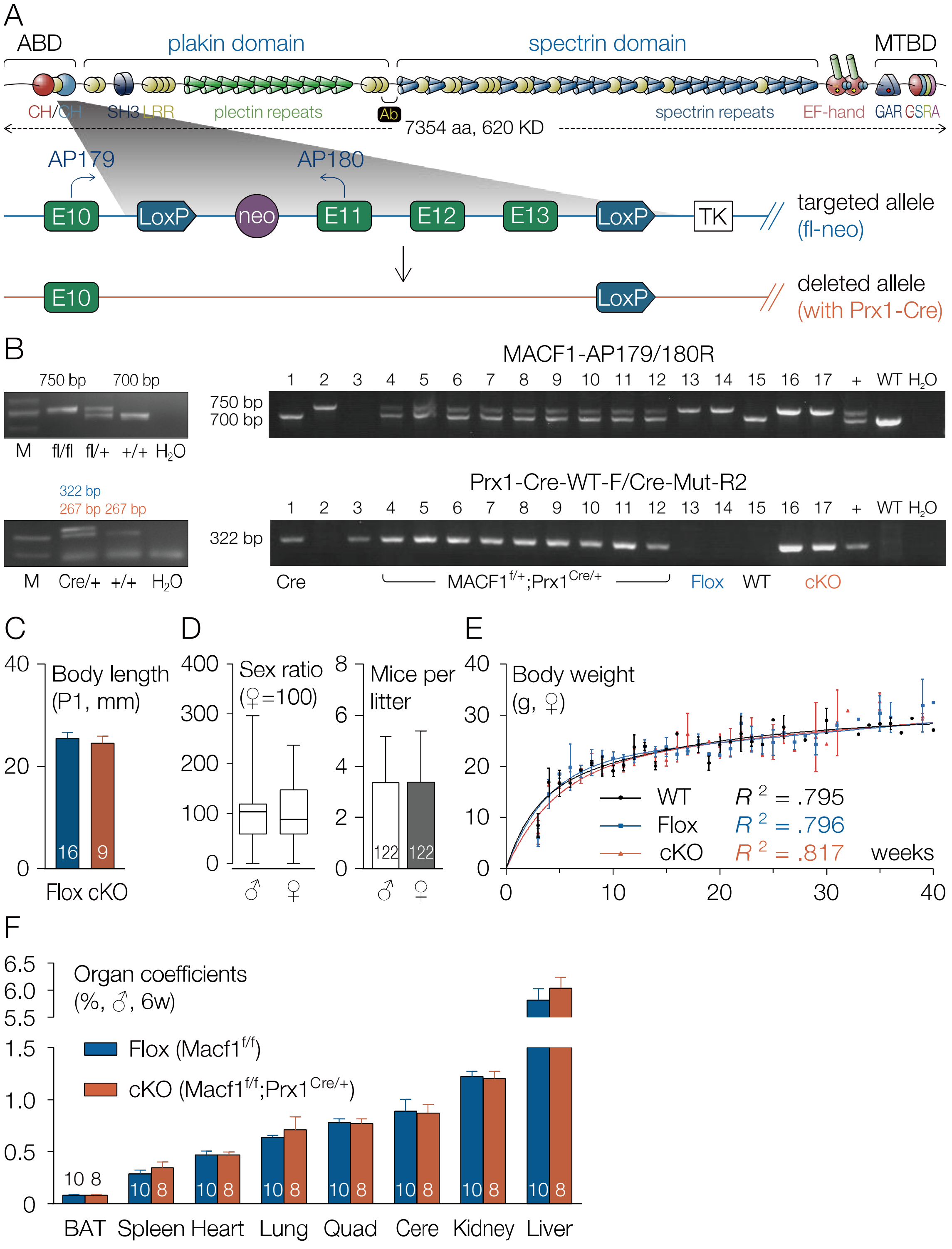


Figure S1. Related to Figure 1.

Fig. S1. Mesenchymal deletion of MACF1 do not apparently affect hereditary and physiological characters in neonatal mice. (A) Schematic diagram showing domain structure of the mouse *Macf1* gene (UniProtKB: Q9QXZ0-1) and targeting strategy for generating the MACF1 conditional knockout mice. The EF1-EF2 domain coordinates two $Ca^{2+}$ ions, while the GAR domain coordinates $Zn^{2+}$. CH, calponin homology; SH3, the SRC homology 3 domain; LRR, leucine-rich repeat; GAR, gas2 (growth arrest specific 2)-related domain; GSRA, tandem repeats of Gly/Ser/Arg/Ala; ABD, actin binding domain. MTBD, microtubule binding domain. Ab, recognition site of the anti-MACF1 antibody (abcam, ab117418). E10, exon 10. Neo, neomycin resistance cassette. TK, thymidine kinase cassette. AP179/180, arbitrary primers designed for genotyping. Prx1-Cre, paired related homeobox 1 promoter driven Cre recombinase expressing mouse. (B) PCR bands for genotyping. (C) Body length of neonatal mice at P1. Body length was defined as ventral distance between the mouth and anus. (D) Sex ratio and litter size in the offspring. Data were collected within 122 litters. (E) Growth curve of female MACF1 cKO mice. Least-squares fitting Gompertz function was used. (F) Organ coefficients of 6-week old male mice. Data were normalized by body weight. BAT, brown adipose tissue. Quad, quadriceps. Cere, cerebrum. Data are represented as mean ± s.d.

### Figure S2

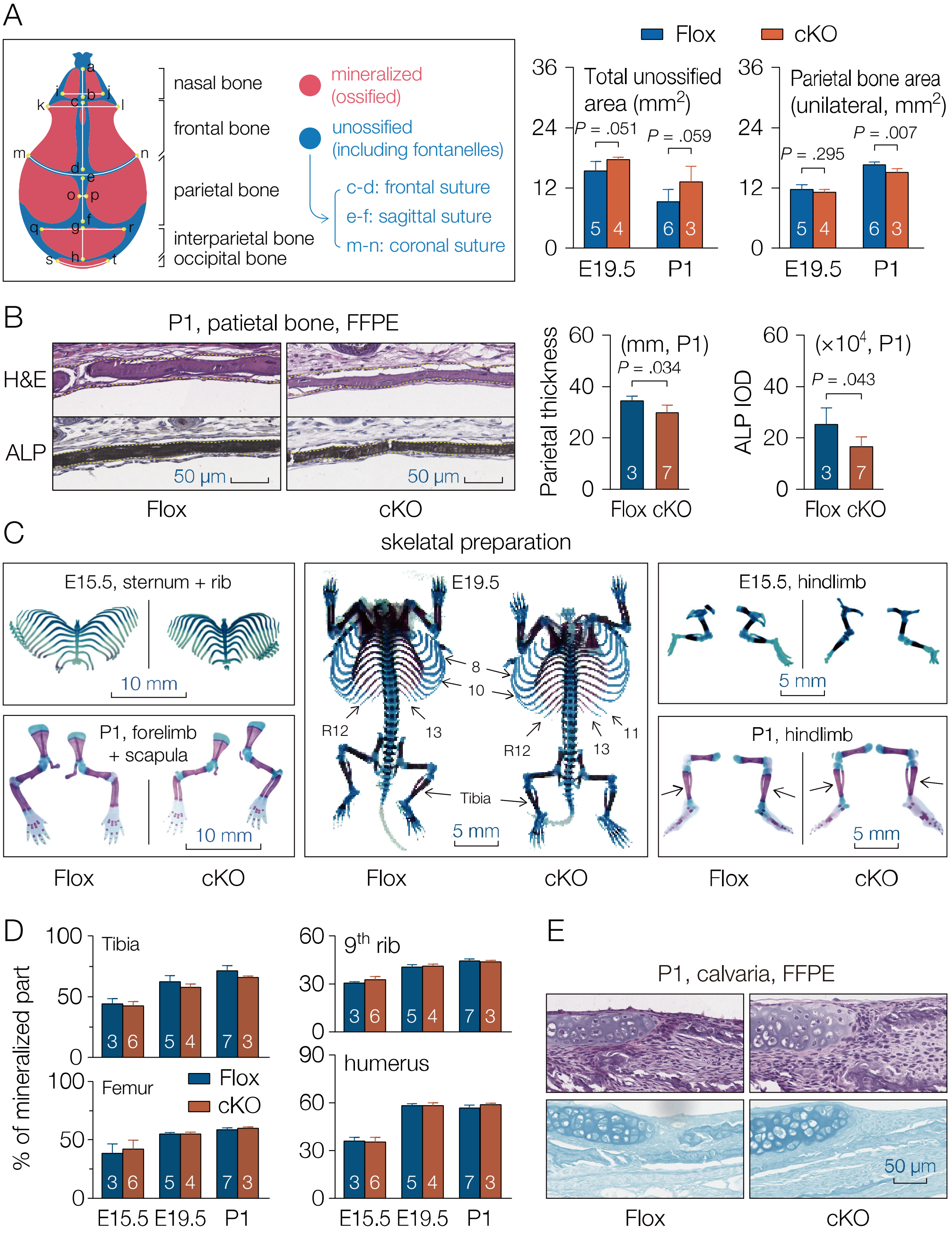


Figure S2. Related to Figure 1.

Fig. S2. Mesenchymal deletion of MACF1 retards early stage bone development in mice. (A) Morphometric analysis of the skull in neonatal mice. A schematic graph showing structure and morphology of neonatal mouse is provided. (B) Representative images of hematoxylin and eosin (H&E) and alkaline phosphatase (ALP) staining showing morphology and ALP expression in the parietal bone (P1, coronal sections). (C) Representative images of cleared skeletal preparation showing ossification in long bones. R12, 12th rib. (D) Ratio of mineralized region in 9th rib, humerus, tibia and femur. (E) Representative images of alcian blue staining showing morphology of chondrocyte in calvaria. Data are represented as mean ± s.d , and compared with littermate controls. Statistical significances were determined using student's *t*-test.

Figure S3

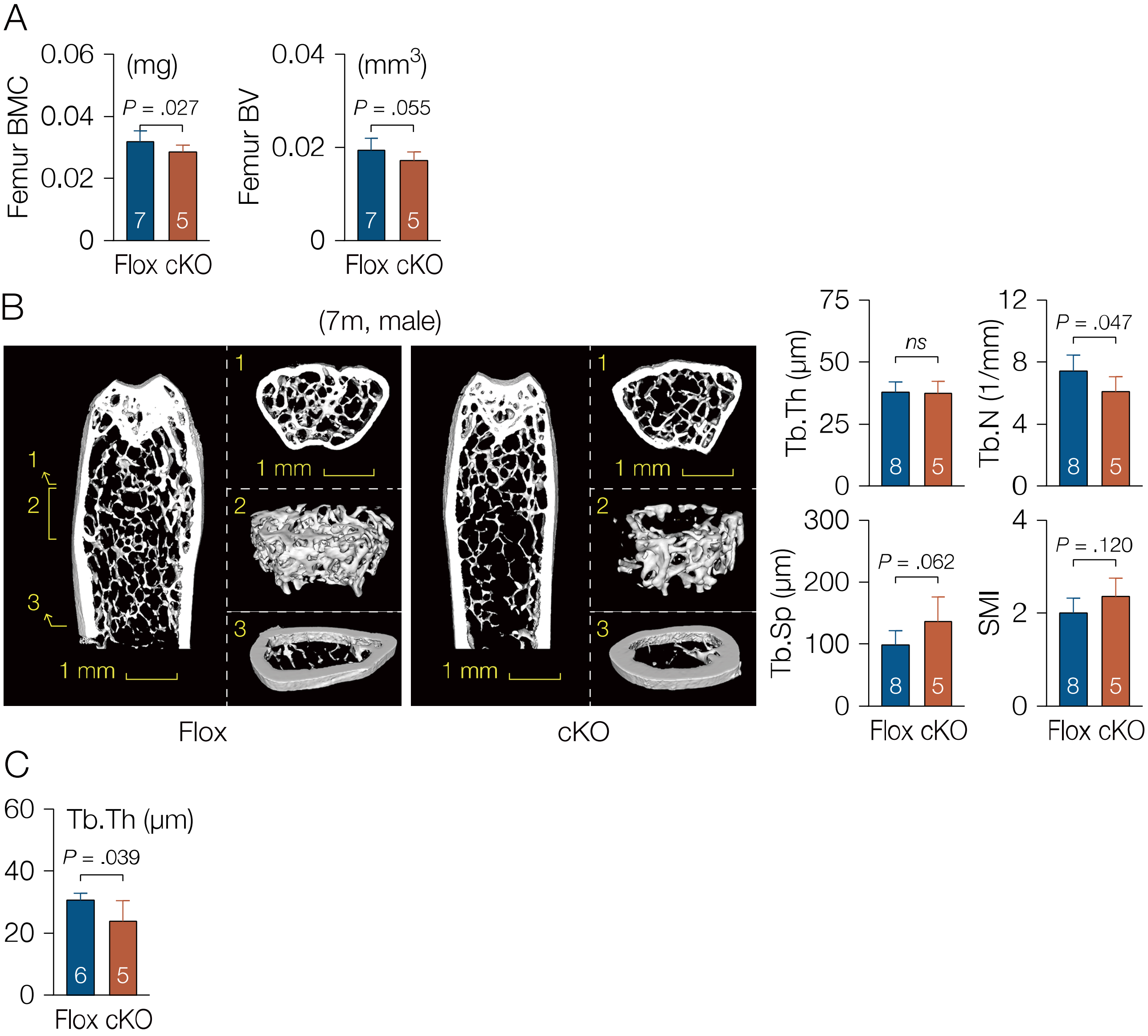


Figure S3. Related to Figure 2.

Fig. S3. Mesenchymal deletion of MACF1 weakens bone properties in adult mice. (A) Mineral content and density of femur and 3rd lumbar vertebra (LV3) in 3-month old male mice.

(B) Representative 3D reconstruction images showing microarchitecture of distial femur (male, 7-month old). Positions of reconstructed region are indicated by arabic numerals. Stereological parameters for trabecular and cortical bone in distal femur are provided. (C) Trabecular thickness in distal femur of 12-month old mice. Data are represented as mean ± s.d. Statistical significance were determined using student's *t*-test.

#### Figure S4

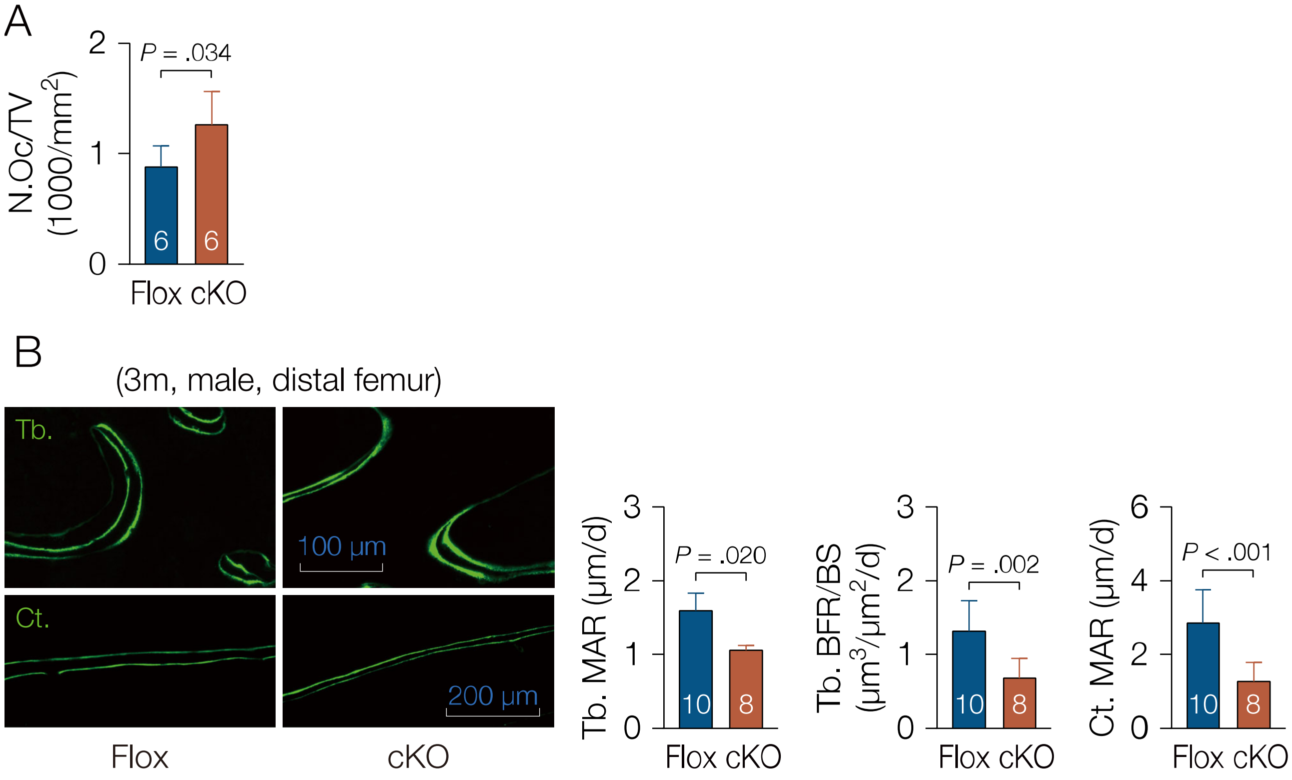


Figure S4. Related to Figure 3.

Fig. S4. Low bone formation capacity in MACF1 cKO mice. (A) Number of osteoclasts per tissue volume in femur of 6-week old male mice. (B) Representative images of calcein double labeling showing mineral apposition and bone formation in femur (male, 3-month old, coronal sections). Tb., trabecular; Ct., cortical. Data are represented as mean ± s.d. Statistical significance were determined using student's *t*-test.

### Figure S5

A

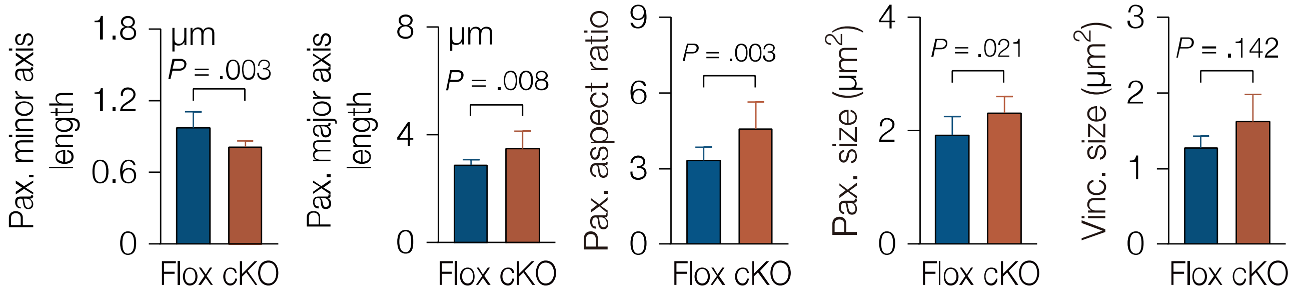


Figure S5. Related to Figure 4.

Fig. S5. (A) Size and morphometric parameters of paxillin or vinculin in MACF1 deficient MSCs. Data are represented as mean ± s.d. Statistical significance were determined using student's *t*-test.

### Figure S6

A

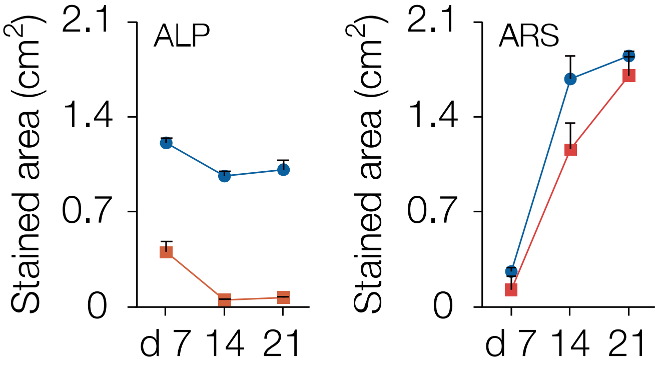


Figure S6. Related to Figure 5.

Fig. S6. Loss of MACF1 inhibits osteoblastic differentiation in MSCs. (A) Stained area of ALP staining and alizarin red staining in osteo-induced MSCs. (B) Immunoblotting analysis of SMAD7, OSTERIX and RUNX2 in osteo-induced MSCs. Data are represented as mean ± s.d.

### Figure S7

A

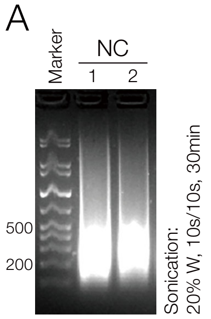


B

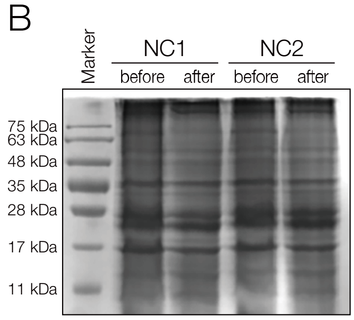


C

| Input gene | Gene ID | Input gene | Gene ID | Input gene | Gene ID |
|---|---|---|---|---|---|
| Fn1 | 2335 | Pdgfa | 5154 | Vcam1 | 7412 |
| Gdf10 | 2662 | Phex | 5251 | Vdr | 7421 |
| Gdf11 | 10220 | Runx2 | 860 | Vegfa | 7422 |
| GDF2 | 2658 | Serpinh1 | 871 | Vegfb | 7423 |
| GDF5 | 8200 | Smad1 | 4086 | Wnt1 | 7471 |
| GDF6 | 392255 | Smad2 | 4087 | Wnt10A | 80326 |
| GDF7 | 151449 | Smad3 | 4088 | Wnt10B | 7480 |
| Gli1 | 2735 | Smad4 | 4089 | Wnt11 | 7481 |
| GSK3A | 2931 | Smad5 | 4090 | Wnt16 | 51384 |
| GSK3B | 2932 | Smad6 | 4091 | Wnt2 | 7472 |
| Icam1 | 3383 | Smad7 | 4092 | Wnt2B | 7482 |
| Igf1 | 3479 | Smad9 | 4093 | Wnt3 | 7473 |
| Igf1r | 3480 | SMURF1 | 57154 | Wnt3A | 89780 |
| IGF2 | 3481 | SMURF2 | 64750 | Wnt4 | 54361 |
| Ihh | 3549 | Sost | 50964 | Wnt5A | 7474 |
| ITGA1 | 3672 | Sox9 | 6662 | Wnt5B | 81029 |
| Itga2 | 3673 | Sp7 | 121340 | Wnt6 | 7475 |
| Itga2b | 3674 | Spp1 | 6696 | Wnt7A | 7476 |
| Itga3 | 3675 | Tgfb1 | 7040 | Wnt7B | 7477 |
| Itgam | 3684 | Tgfb2 | 7042 | Wnt8A | 7478 |
| Itgav | 3685 | Tgfb3 | 7043 | Wnt8B | 7479 |
| Itgb1 | 3688 | Tgfbr1 | 7046 | Wnt9A | 7483 |
| Mmp10 | 4319 | Tgfbr2 | 7048 | Wnt9B | 7484 |
| Mmp2 | 4313 | Tgfbr3 | 7049 | | |
| Mmp8 | 4317 | Tnf | 7124 | | |
| Mmp9 | 4318 | TNFRSF11B | 4982 | | |
| Nfkb1 | 4790 | Tnfsf11 | 8600 | | |
| Nog | 9241 | Twist1 | 7291 | | |

| Input gene | Gene ID | Input gene | Gene ID | Input gene | Gene ID | Input gene | Gene ID |
|---|---|---|---|---|---|---|---|
| Acvr1 | 90 | Bmp5 | 653 | Chrd | 8646 | Csf2 | 1437 |
| Ahsg | 197 | Bmp6 | 654 | Col10a1 | 1300 | Csf3 | 1440 |
| Alpl | 249 | Bmp7 | 655 | Col14a1 | 7373 | CTNNB1 | 1499 |
| Anxa5 | 308 | BMP8a | 353500 | COL15A1 | 1306 | Ctsk | 1513 |
| Bglap | 632 | BMP8b | 656 | Col1a1 | 1277 | Dlx5 | 1749 |
| Bgn | 633 | Bmpr1a | 657 | Col1a2 | 1278 | Egf | 1950 |
| Bmp1 | 649 | Bmpr1b | 658 | Col2a1 | 1280 | EGFR | 1956 |
| BMP10 | 27302 | Bmpr2 | 659 | Col3a1 | 1281 | Fgf1 | 2246 |
| BMP15 | 9210 | CALCR | 799 | Col4a1 | 1282 | Fgf2 | 2247 |
| Bmp2 | 650 | Cd36 | 948 | Col5a1 | 1289 | Fgfr1 | 2260 |
| Bmp3 | 651 | Cdh11 | 1009 | Comp | 1311 | Fgfr2 | 2263 |
| Bmp4 | 652 | Cebpa | 1050 | Csf1 | 1435 | Flt1 | 2321 |

D

| Rank | Gene Symbol | Gene ID | GO: Molecular Function | | GO: Biological Process | | GO: Cellular Component | | Human Phenotype | | Mouse Phenotype | | Pathway | | Pubmed | | Disease | | Average Score | Overall pValue | Interactant count | Score |
|---|---|---|---|---|---|---|---|---|---|---|---|---|---|---|---|---|---|---|---|---|---|---|
| | | | Score | P value | Score | P value | Score | P value | Score | P value | Score | P value | Score | P value | Score | P value | Score | P value | | | | |
| 1 | SMAD7 | 4092 | 1 | .00039 | 1 | .09005 | .99018 | .01665 | .96196 | .03815 | 1 | .04822 | 1 | .00407 | 1 | .03970 | 1 | .03195 | .99402 | .00007 | 104 | .00066 |
| 2 | REL | 5966 | .99753 | .01123 | 1 | .09005 | .78245 | .08443 | N/A | 0 | 1 | .04822 | .88625 | .02033 | 1 | .03970 | 1 | .03486 | .96037 | .00179 | 224 | .00024 |
| 3 | HES1 | 3280 | .99616 | .01278 | 1 | .09005 | .53917 | .13749 | N/A | 0 | 1 | .04822 | .67410 | .03311 | 1 | .03970 | 1 | .03195 | .90833 | .00268 | 52 | .00009 |
| 4 | BHLHE40 | 8553 | .99966 | .00445 | 1 | .09682 | .73668 | .09624 | N/A | 0 | 1 | .04822 | .30427 | .06158 | 1 | .03970 | .97084 | .07591 | .89620 | .00285 | 28 | .00006 |
| 5 | ZFP748 | 212276 | DATA NOT FOUND | | | | | | | | | | | | | | | | | | | |

E

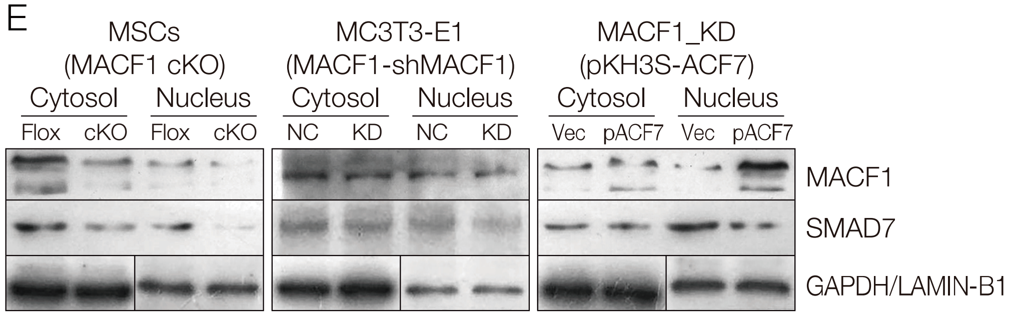


Figure S7. Related to Figure 5.

Fig. S7. MACF1 interacts with SMAD7. (A) Gel electrophoresis for analysis of DNA quality in supernatant of MC3T3-E1 lysate after sonication. (B) Coomassie blue staining for determination of protein quality in supernatant of MC3T3-E1 lysate before and after sonication. (C) 127 osteogenic differentiation-related genes entered into the ToppGene database as training set. (D) Summary of output results predicted by the ToppGene database. The *Score* value represents how much related a queried gene is with osteogenic differentiation. (E) Western blot analysis of MACF1 and SMAD7 levels in MACF1 cKO MSCs, MACF1 knockdown preosteoblasts, and MACF1 stable transfected MACF1 knockdown preosteoblasts. ACF7 (Actin cross-linking family protein 7) is a synonym for MACF1. Lamin B1 and GAPDH were used as internal reference for nucleus and cytoplasm, respectively.

### Figure S8

A

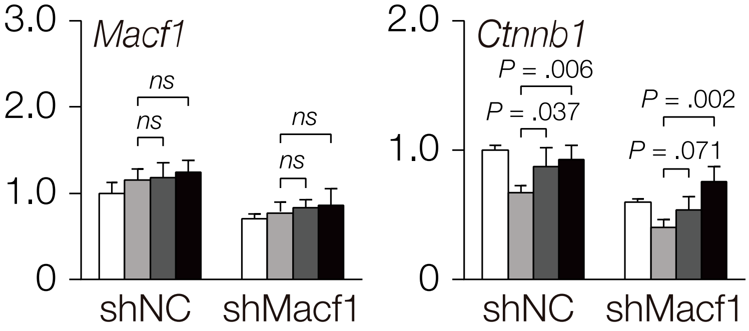


Figure S8. Related to Figure 7.

Fig. S8. MACF1 facilitate SMAD7 nucleus translocation to drive osteoblastic differentiation. (A). Real-time PCR analysis of *Macf1* and *Ctnnb1* relative gene expression in Macf1 knockdown MC3T3-E1 preosteoblasts after plasmid transfection.
